# Somatic mosaicism in amyotrophic lateral sclerosis and frontotemporal dementia identifies focal mutations associated with widespread degeneration

**DOI:** 10.1101/2023.11.30.569436

**Authors:** Zinan Zhou, Junho Kim, August Yue Huang, Matthew Nolan, Junseok Park, Ryan Doan, Taehwan Shin, Michael B. Miller, Mingyun Bae, Boxun Zhao, Jinhyeong Kim, Brian Chhouk, Katherine Morillo, Rebecca C. Yeh, Connor Kenny, Jennifer E. Neil, Chao-Zong Lee, Takuya Ohkubo, John Ravits, Olaf Ansorge, Lyle W. Ostrow, Clotilde Lagier-Tourenne, Eunjung Alice Lee, Christopher A. Walsh

**Affiliations:** Division of Genetics and Genomics, Boston Children’s Hospital, Boston, MA, USA; Manton Center for Orphan Disease, Boston Children’s Hospital, Boston, MA, USA; Department of Pediatrics, Harvard Medical School, Boston, MA, USA; Department of Biological Sciences, Sungkyunkwan University, Suwon, South Korea; Department of Neurology, The Sean M. Healey and AMG Center for ALS at Mass General, Massachusetts General Hospital, Harvard Medical School, Boston, MA, USA; Department of Pathology, Brigham and Women’s Hospital, Harvard Medical School, Boston, MA, USA; Howard Hughes Medical Institute, Boston Children’s Hospital, Boston, MA, USA; Department of Neurology, Yokohama City Minato Red Cross Hospital, Yokohama, Kanagawa, Japan; Department of Neurosciences, School of Medicine, University of California San Diego, La Jolla, CA, USA; Nuffield Department of Clinical Neurosciences, University of Oxford, Oxford, Oxfordshire, UK; Department of Neurology, Lewis Katz School of Medicine at Temple University, Philadelphia, PA, USA

## Abstract

Although mutations in many genes cause familial amyotrophic lateral sclerosis and frontotemporal dementia, most cases are sporadic (sALS and sFTD) with unclear etiology. We tested whether somatic mutations contribute to sALS and sFTD by deep targeted sequencing of 88 neurodegeneration-related genes in postmortem brain and spinal cord samples from 399 sporadic cases and 144 controls. Predicted deleterious somatic variants in ALS/FTD genes were observed in 2.1% of sporadic cases lacking deleterious germline variants. These variants occurred at very low allele fractions (typically <2%) and were often focal and enriched in disease-affected regions. Analysis of bulk RNA-seq data from an additional cohort identified deleterious somatic variants in *DYNC1H1* and *LMNA*, genes associated with pediatric motor neuron degeneration. Targeted long-read sequencing further identified one sFTD case with *de novo* somatic *C9orf72* repeat expansions. Together, these findings suggest that rare, focal somatic variants can contribute to sALS and sFTD and drive widespread neurodegeneration.

## Introduction

Amyotrophic lateral sclerosis (ALS), a disease in which premature loss of upper and lower motor neurons (UMNs and LMNs) leads to fatal paralysis, shows clinical, genetic, and pathological overlap with frontotemporal dementia (FTD), a neurodegenerative disorder characterized by behavioral, language, and memory dysfunction^1^. 5-22% of individuals with ALS develop FTD, and ∼15% of those with FTD eventually develop ALS^2^. ALS and FTD also share common pathology, with cytoplasmic inclusions of TAR DNA binding protein (TDP-43) found in almost all ALS brains and in half of FTD brains^3,4^. ALS typically begins focally and spreads regionally as the disease progresses^5,6^, although whether degeneration begins in UMNs, LMNs, or both simultaneously has remained controversial^7,8^, with some studies suggesting that focality can manifest independently in UMNs and LMNs^5,9^. TDP-43 pathology also follows stereotypical patterns in ALS and FTD brains^9–11^, thought to reflect focal onset and intercellular transmission of TDP-43 inclusions in a prion-like manner, as shown in cell and animal models^12–18^.

Whereas over 30 genes are implicated in ALS and FTD^19^, most causative genes are linked to familial ALS (fALS) and FTD (fFTD), while 90-95% of cases are sporadic ALS (sALS) and FTD (sFTD) without a family history^20^. The focal onset of ALS and FTD, their stereotypical spread, and the increased risk in smokers^21^ have raised interest in potential roles of somatic mosaic mutations in the pathogenesis of ALS and FTD^22^. Somatic mutations are increasingly recognized as prevalent in normal-appearing tissues, but somatic mutations responsible for neurological conditions are often limited to the central nervous system (CNS)^23^, and hence undetectable through DNA sequencing of non-CNS tissues. Recent studies have evaluated the contributions of somatic mutation to Alzheimer’s and Parkinson’s disease directly using postmortem brain tissues^24^.

In this study, we assessed potential contributions of somatic variants—distinguished by their variant allele frequencies (VAFs)—to sALS and sFTD using deep sequencing of a panel of neurodegeneration/dementia-associated genes on postmortem tissues of various brain regions and spinal cords from 399 unique sALS and sFTD cases. Our study identified novel predicted deleterious somatic variants in 2.1% of the sALS and sFTD cases without pathogenic or predicted deleterious germline variants. Protein-altering (missense/nonsense/frameshift) somatic variants showed enrichment in sALS and sFTD cases and in disease-affected brain regions, supporting roles in disease pathogenesis. Regional analysis revealed focality of predicted deleterious somatic variants in primary motor cortex and spinal cord, supporting independent disease initiation in UMNs and LMNs, but also strongly supporting models of ALS and FTD in which the disease spreads beyond a relatively confined region containing a somatic variant. Complementary analyses of bulk RNA-seq and targeted long-read sequencing of brain and spinal cord tissues further revealed somatic variants in genes not previously associated with ALS or FTD, as well as a *de novo* somatic *C9orf72* repeat expansion. Together, our study opens new avenues for understanding the etiology of sporadic ALS and FTD.

## Results

### Deep targeted sequencing of neurodegenerative genes in sALS and sFTD brains

To directly detect somatic variants in sALS and sFTD brains, we obtained post-mortem frozen tissues of several brain regions and spinal cords from individuals diagnosed with sALS or sFTD, as well as from age-matched controls through the Massachusetts Alzheimer’s Disease Research Center, Oxford Brain Bank, and Target ALS Foundation (Fig. 1a and Supplementary Table 1). Additional brain tissues from ALS, FTD and control cases, without a record of family history but with an age of death above 45 years old, were also obtained from the NIH NeuroBioBank. We performed molecular inversion probe (MIP)-panel sequencing^25^ of 88 neurodegeneration-associated genes at ∼1,800× deduplicated depth across 1,787 samples from 291 ALS, 117 FTD, and 144 neurotypical control individuals (Fig. 1a,b, Supplementary Fig. 1, Supplementary Table 2, and Supplementary Note).

**Fig. 1.**
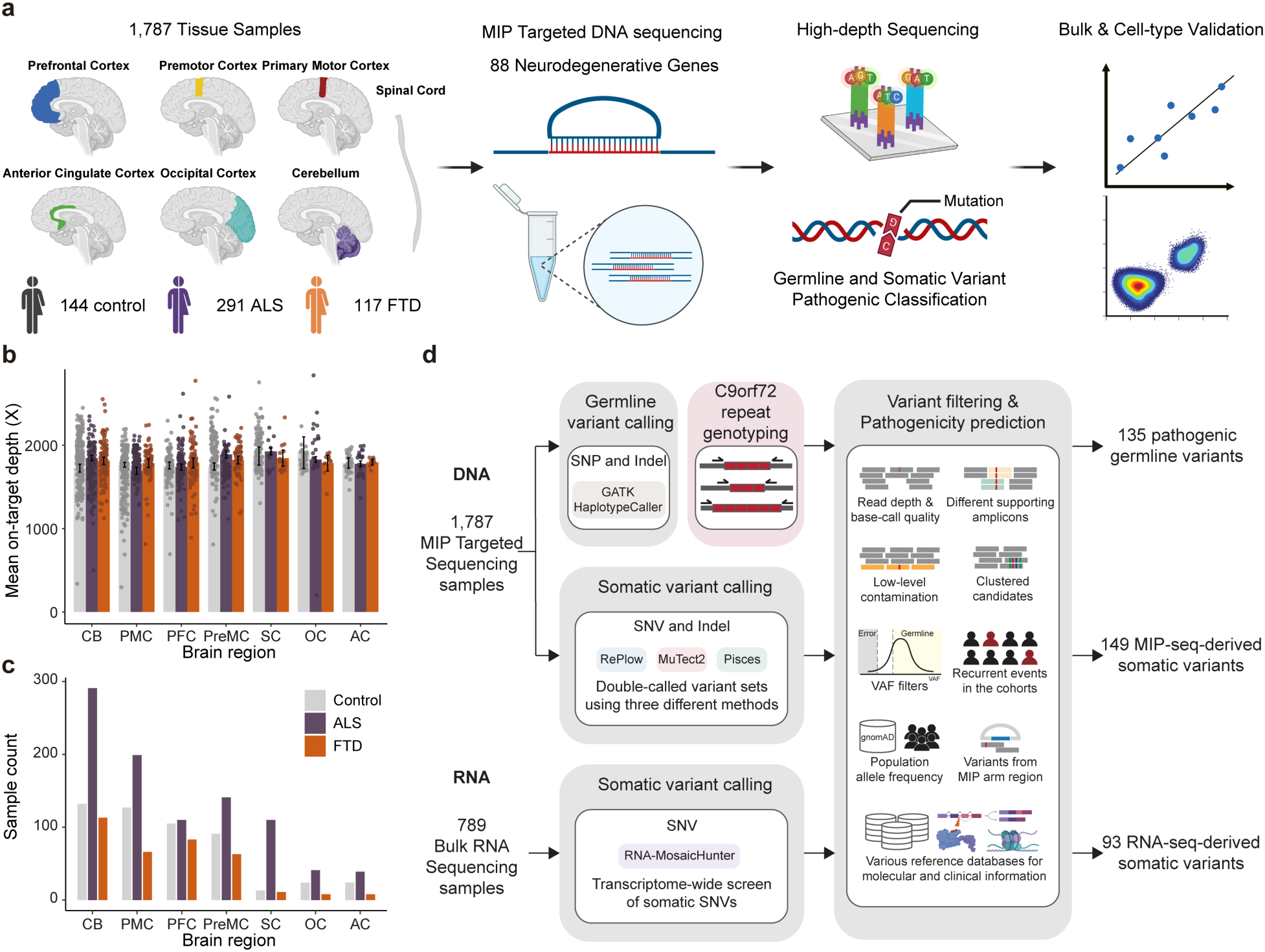
Experimental and analysis strategies. **a**, Overall scheme of the experiments. Genomic DNA isolated from 1,787 postmortem tissue samples of multiple brain regions and spinal cords of 144 control, 291 ALS, and 117 FTD cases were used for molecular inversion probe (MIP) capture sequencing with ultra-high depth. Created with BioRender.com. **b**,**c**, Mean sequencing depth (**b**) and number of tissue samples (**c**) in different brain regions and spinal cords of control, ALS, and FTD cases. Control, *n* = 516; ALS, *n* = 938; FTD, *n* = 375. Note that 42 samples from 9 ALS-FTD cases were included in both conditions. CB, cerebellum; PMC, primary motor cortex; PFC, prefrontal cortex; PreMC, premotor cortex; SC, spinal cord; OC, occipital cortex; AC, anterior cingulate cortex. Error bars, 95% CI. **d**, Methodological pipelines to identify germline and somatic variants. Germline variants were called by GATK HaplotypeCaller. *C9orf72* genotype of ALS and FTD cases were determined by repeat-primed PCR. Somatic variants were called by RePlow, MuTect2, and Pisces. Additional somatic variants were called from 789 bulk RNA-seq profiles of multiple brain regions and spinal cords of ALS cases generated by the New York Genome Center ALS Consortium using RNA-MosaicHunter. Created with BioRender.com.

### Pathogenic germline variants in sALS and sFTD cases

We first identified pathogenic germline single-nucleotide variants (SNVs) and short insertions and deletions (indels) using GATK followed by stringent filtering (Fig. 1d), with functional annotation and deleteriousness prediction performed using ANNOVAR^26^ and multiple clinical databases. In addition, *C9orf72* repeat expansions, the most common inherited cause of ALS and FTD^27,28^, were genotyped by a repeat-primed PCR assay (Supplementary Fig. 2). Overall, 20.6% (60/291) of ALS, 25.6% (30/117) of FTD and 0.7% (1/144) of control cases carried *C9orf72* repeat expansions or pathogenic germline mutations in ALS and FTD genes (Fig. 2a and Supplementary Tables 3 and 4). Missense variants represented the most prevalent variant type (Fig. 2b). *C9orf72* was the most frequently mutated gene, followed by *SOD1* in ALS, and *GRN* and *MAPT* in FTD (Fig. 2c,d). The overall fractions of *C9orf72* repeat expansion carriers (10.6% in ALS; 12.0% in FTD) were slightly higher than previously reported in sporadic cases, but still lower than in familial cases^29–31^.

**Fig. 2.**
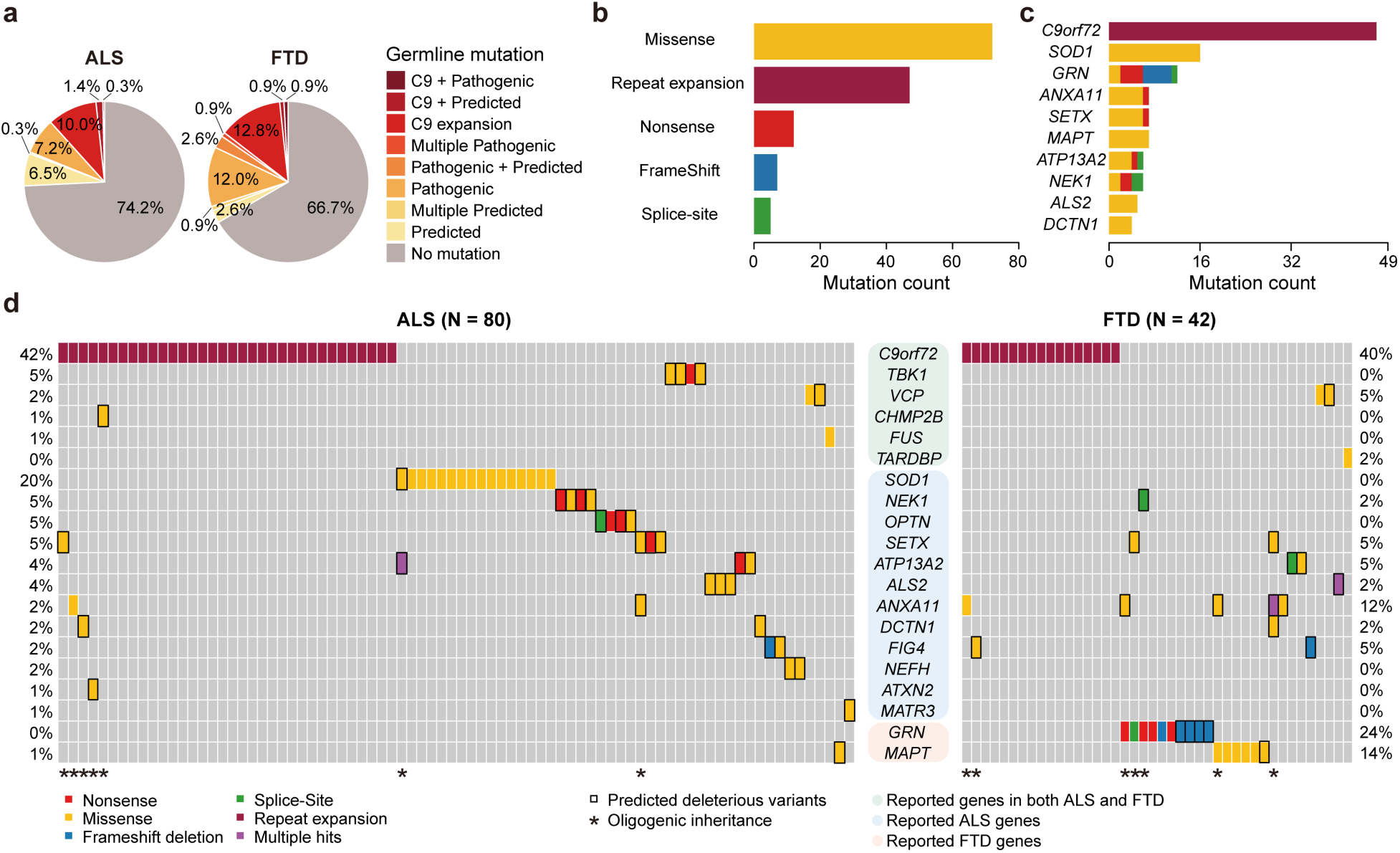
C*9*orf72 repeat expansion and pathogenic germline variants in ALS/FTD genes are prevalent in ALS and FTD. **a**, Proportions of ALS and FTD cases with *C9orf72* repeat expansion, pathogenic, and predicted deleterious germline variants in ALS/FTD genes. Cases with multiple predicted deleterious variants are indicated with ‘+’ sign. Note that cases with a heterozygous variant in recessive genes (e.g., *ALS2*, *ATP13A2*) are not included in the proportions. **b**, Distribution of *C9orf72* repeat expansion, pathogenic, and predicted deleterious germline variants in ALS/FTD genes classified by variant types. **c**, Ranking of the top 10 mutated ALS/FTD genes. **d**, Visualization of ALS and FTD cases (vertical columns) with pathogenic and predicted deleterious germline variants (horizontal rows) in ALS/FTD genes. Color codes indicate the types of variants. Rectangular outline represents predicted deleterious variants. Genes are grouped by their known involvement in the diseases. * indicates cases with multiple pathogenic or predicted deleterious variants.

In addition, predicted deleterious germline variants in dominant ALS/FTD genes were found in an additional 14.1% of ALS, 18.8% of FTD, and 4.9% of control cases with significant enrichment observed in both ALS (OR = 3.20, 95% CI: 1.37-8.69, *P* = 3.2 × 10^-3^) and FTD (OR = 4.51, 95% CI: 1.77-13.01, *P* = 5.5 × 10^-4^) cases (Fig. 2a and Supplementary Tables 3 and 4). While most predicted deleterious variants were non-synonymous SNVs requiring functional validation, two novel *GRN* frameshift variants (p.L46Rfs*18 and p.D250Tfs*6) identified in FTD cases were probably pathogenic (Supplementary Table 3), since loss-of-function *GRN* mutations are known to cause FTD in a dominant manner^32,33^. Consistent with previous reports, we found evidence of possible oligogenic inheritance. This includes one case carrying both *C9orf72* expansion and a pathogenic variant in *ANXA11*, as well as 12 additional cases carrying combinations of pathogenic and predicted deleterious variants (Fig. 2d and Supplementary Table 3)^34–37^. Interestingly, we also identified cross-disease—variants in ALS genes in FTD and vice versa—highlighting potential shared genetic mechanisms (Fig. 2d).

### Identification of somatic SNVs and indels from MIP sequencing data

We developed a custom pipeline integrating RePlow^38^, Mutect2^39^, and Pisces^40^ for calling somatic SNVs and indels in our MIP sequencing data (Fig. 1d). We selected somatic variants identified by at least two of the three callers (double-called variants) followed by multi-step variant filters to remove false positive candidates. Unlike heterozygous germline variants with VAFs around 50%, heterozygous somatic variants have VAFs less than 50%, and we only called somatic variants with VAFs below 35%. To benchmark our pipeline, we performed spike-in experiments by mixing two human samples from the Genome in a Bottle Consortium (GIAB) at low VAFs and observed high sensitivity and precision with a low false positive rate across VAFs (Extended Data Fig. 1a,b and Supplementary Note).

We applied our custom pipeline to 1,787 samples to obtain initial somatic variant candidates. Among these, we identified and removed candidates arising from potential sample contamination in 29 samples (Supplementary Note). The remaining call set contained 98 unique somatic SNV and indel candidates (a total of 149 candidates, including shared ones observed in multiple regions of the same individuals; see Supplementary Table 5). Variants with low VAFs (<5%) were more common in disease cases than in normal controls (Extended Data Fig. 2). All somatic candidates were validated by deep amplicon sequencing, except one for which primers could not be designed; 34 exonic candidates were additionally assessed by droplet digital PCR (ddPCR). Thirteen candidates were actually germline variants, consistent with known VAF deviation in MIPs sequencing due to probe hybridization and amplicon design biases^41,42^. After excluding these, 64 out of 85 candidates (75.2%) were validated as somatic (Supplementary Table 6). The VAFs of validated candidates in amplicon sequencing showed a strong correlation with their original VAFs in the MIP sequencing data (Fig. 3a). All subsequent somatic variant analyses were conducted using only the validated candidates.

**Fig. 3.**
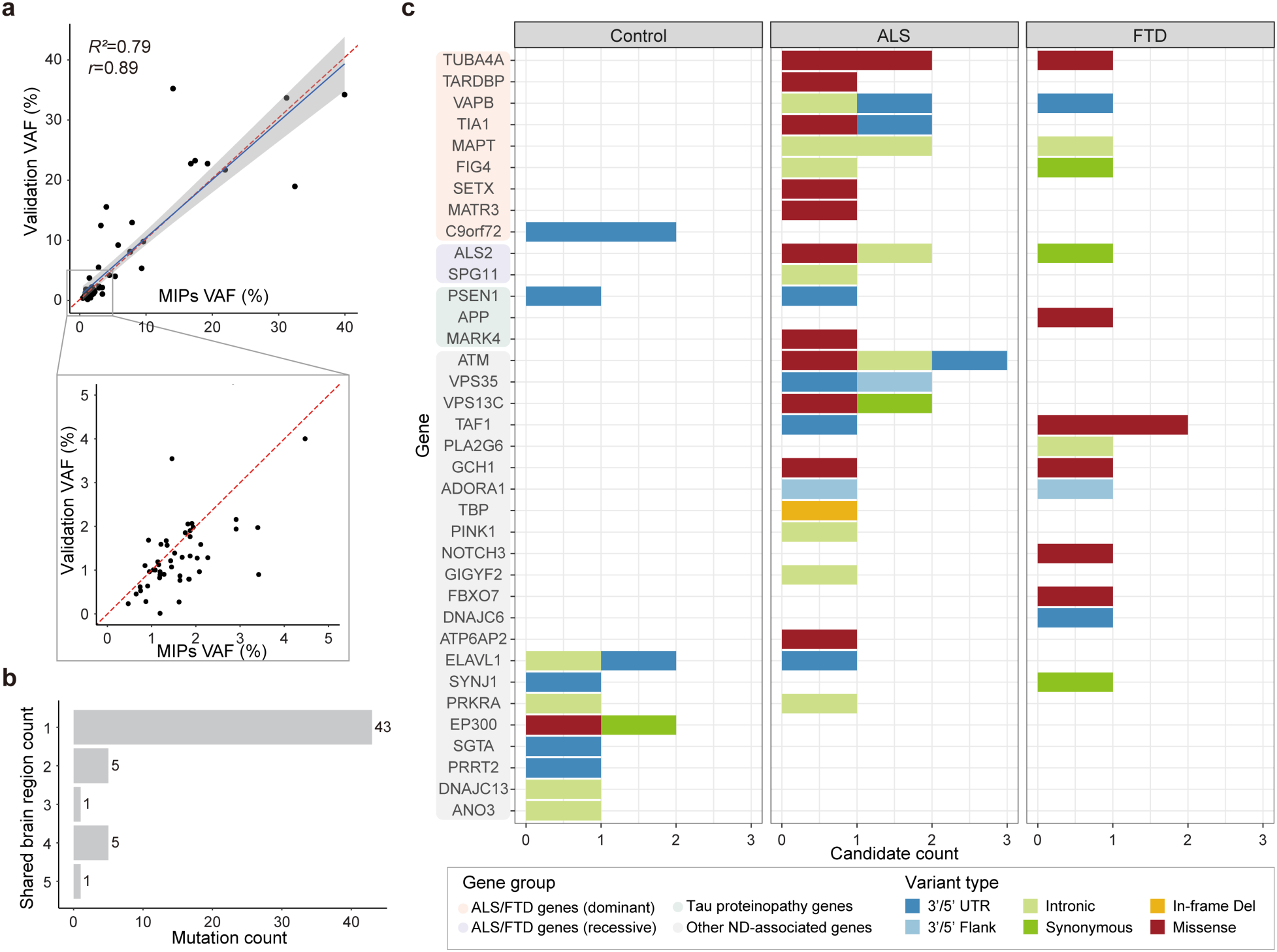
Somatic variants in MIP sequencing data tend to be focal, protein-altering, and are almost exclusively restricted to disease cases. **a**, The observed VAFs of somatic variants in amplicon sequencing validation were consistent with the VAFs in original MIP sequencing. Sixty-four somatic variants were validated and included in the plot. Shaded band, 95% CI of the fitted linear regression. **b**, Total somatic variant counts classified by the number of brain regions in which a given variant was identified. **c**, Distribution of somatic variants in all neurodegenerative genes. Genes are categorized into four groups: ALS/FTD-related dominant genes, ALS/FTD-related recessive genes, Tau proteinopathy-related genes, and other neurodegeneration/dementia-associated genes. Color codes indicate variant types. Note that somatic variants identified in controls are unlikely to alter function, with just one missense variant (red) and the remaining being synonymous or noncoding substitutions.

### Somatic variants in disease-relevant genes are enriched in ALS and FTD cases lacking pathogenic germline variants

To examine the burden and potential roles of somatic variants in ALS and FTD, we focused on cases lacking pathogenic or predicted deleterious germline variants in ALS/FTD genes (germline-free cases), including cases with heterozygous germline variants in recessive ALS/FTD genes (e.g., *ATP13A2*, *ALS2*) that were predicted to be deleterious. Fifty-five unique somatic variants across the targeted regions were identified among 696, 243, and 516 samples from 216 germline-free ALS cases, 78 germline-free FTD cases, and 144 neurotypical controls, respectively. Most of them (78.2%, 43/55) were focal to a single tissue region and present at very low VAFs (Fig. 3b and Extended Data Fig. 2), consistent with late-arising^43^, CNS-restricted events. Mutational signature analysis ^44^ revealed predominance of clock-like signatures (SBS5 and SBS1) (Extended Data Fig. 3), consistent with previous studies of somatic mutagenesis in the normal human brain^45,46^.

Across all targeted neurodegeneration/dementia-related genes, protein-altering somatic variants showed clear separation between disease and control groups (Fig. 3c), with only one such variant observed in controls compared with 15 in ALS and 7 in FTD cases. Both exonic and protein-altering somatic variants were significantly enriched in ALS and FTD (Fig. 4a; exonic: *P* = 0.044 and *P* = 0.0014; protein-altering: *P* = 0.019 and *P* = 0.010; linear mixed model; see Supplementary Note), whereas intronic and non-coding somatic variants showed no enrichment. Consistently, ratios of non-synonymous to non-coding variants were increased in ALS and FTD at both group and individual levels (Extended Data Fig. 4), indicating disease-specific accumulation of potentially damaging variants.

**Fig. 4.**
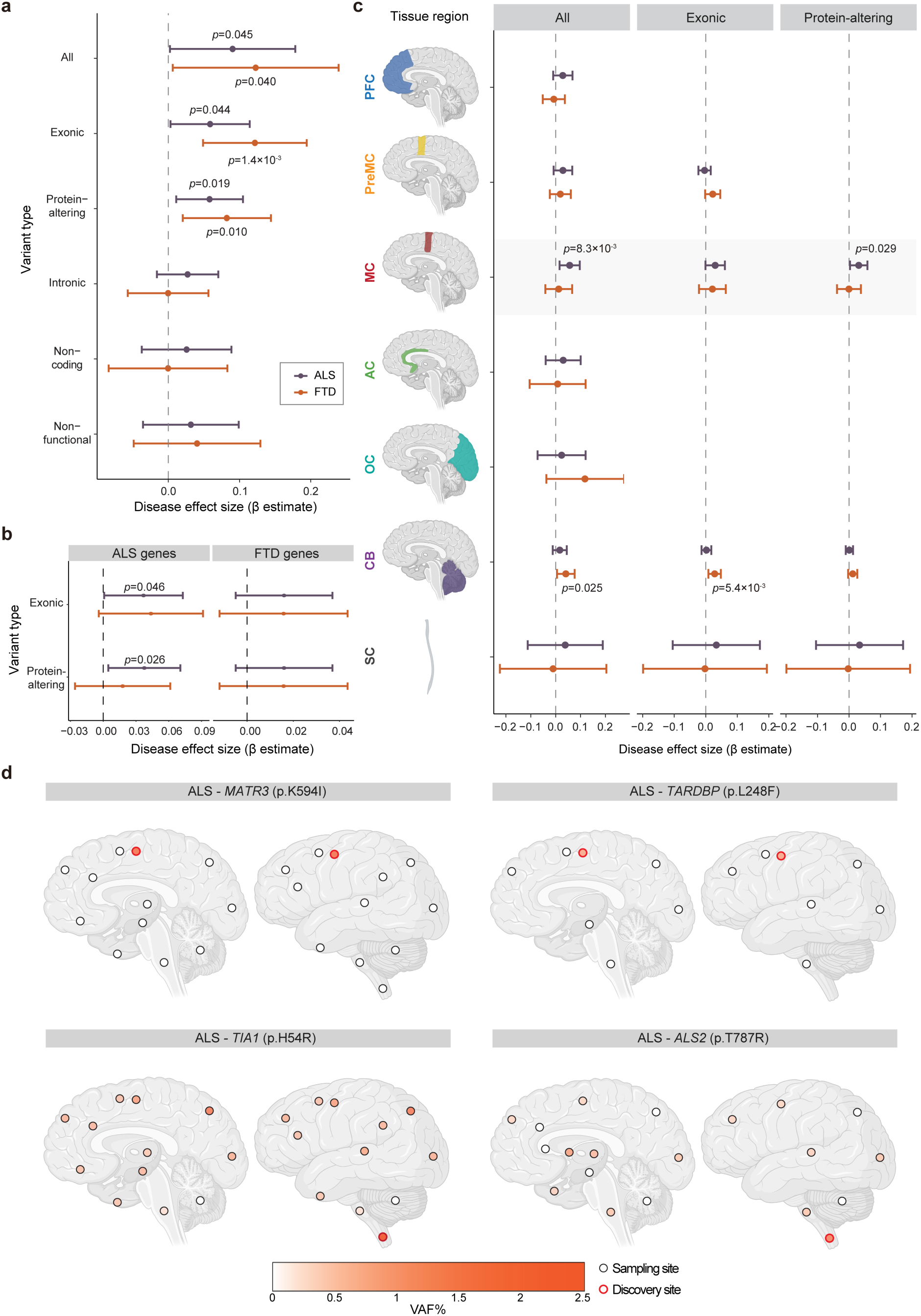
Somatic variants are enriched in ALS and FTD cases and disease-related tissue regions. **a**, Enrichment of somatic variants in different genomic regions of germline-free ALS and FTD cases compared to normal controls. **b**, Enrichment of exonic and protein-altering somatic variants in two different groups of disease-related genes (ALS genes and FTD genes) compared to normal controls. **c**, Enrichment of somatic variants in ALS/FTD genes across different brain regions of germline-free ALS and FTD cases compared to normal controls. Missing points indicate no variants were observed. **a**-**c**, Disease effect sizes were estimated using an individual-wise linear mixed-effects model by comparing variant burden per germline-free disease individual with that of normal individuals (see Methods). Significance and 95% CI were estimated while controlling for potential confounding factors, including average read depth, sex, post-mortem interval, sequencing batch, and number of samples per donor. Sample sizes were: control, *n* = 144; ALS, *n* = 216; FTD, *n* = 78 (biological replicates). Unadjusted *P* values are shown, as the tests examine biologically distinct but non-independent categories with differing background mutation structures. Created with BioRender.com. **d**, Regional distribution of VAFs of several predicted deleterious somatic variants in individual brains and spinal cords. Brain cortex is annotated by Brodmann areas. The color spectrum indicates the VAFs of somatic variants in amplicon sequencing. White indicates regions without the somatic variants. Red highlight indicates the region of initial detection by MIP sequencing. Created with BioRender.com.

When restricted to ALS/FTD genes, exonic and protein-altering variants in established ALS genes were specifically enriched in germline-free ALS cases (Fig. 4b; exonic: *P* = 0.046; protein-altering: *P* = 0.026; linear mixed model). In contrast, germline-free FTD cases showed no enrichment in known FTD genes but exhibited significant enrichment in other neurodegeneration/dementia-related genes (Fig. 4a,b and Extended Data Fig. 5), consistent with broader pathological heterogeneity of FTD, and perhaps implicating a shared genetic architecture with other neurodegenerative diseases. No protein-altering variants in ALS/FTD genes were observed in controls or in ALS/FTD cases carrying pathogenic germline variants (Supplementary Tables 6 and 7).

Somatic variants also exhibited region-specific enrichment in disease-affected regions of germline-free ALS and FTD cases (Fig. 4c and Extended Data Fig. 6). Across all targeted genes, enrichment was observed in the primary motor cortex in ALS cases (Extended Data Fig. 6; all: *P* = 0.020; exonic: *P* = 0.026; protein-altering: *P* = 0.012; linear mixed model) and in the prefrontal cortex of FTD brains (Extended Data Fig. 6; exonic: *P* = 2.3 × 10^-3^; protein-altering: *P* = 7.7 × 10^-4^; linear mixed model). In contrast, the premotor cortex—located immediately between the primary motor cortex and prefrontal cortex—showed no enrichment for either condition. When restricted to ALS/FTD genes, ALS cases still demonstrated the enrichment only in the primary motor cortex (Fig. 4c; all: *P* = 8.3 × 10^-3^; exonic: *P* = 0.060; protein-altering: *P* = 0.029; linear mixed model), while FTD cases showed no significant enrichment in the prefrontal cortex, further supporting a broader genetic heterogeneity in FTD. The spinal cord in ALS showed a modest increase in exonic and protein-altering variants, although this analysis is limited by the small number of control samples and wide confidence intervals (Fig. 4c). Together, these diagnosis- and region-specific patterns suggest that functional somatic variants may contribute to the pathogenesis of sALS and sFTD.

### Predicted deleterious somatic variants have restricted regional distributions and are enriched in hypodiploid cells

Pathogenicity prediction identified six predicted deleterious somatic SNVs in known ALS and FTD genes (Supplementary Table 7), which account for 2.3% and 1.3% of germline-free ALS and FTD cases, respectively (2.1% for overall). All variants in ALS cases were observed in primary motor cortex or spinal cord, the most severely affected regions in ALS, emphasizing remarkable topographic specificity. All somatic variants occurred in disease genes with dominant inheritance when found in the germline setting, except for one sALS case with a predicted deleterious somatic *ALS2* (p.T787R) variant identified in the spinal cord. *ALS2* is an autosomal recessive disease gene^47,48^, and this individual also carried a predicted deleterious germline *ALS2* (p.Q24R) variant in addition to the identified somatic variant. Both *ALS2* variants were predicted to be deleterious, consistent with a “second hit” mechanism at the cellular level in a small proportion of spinal cord cells.

We selected four predicted deleterious somatic SNVs—*TIA1* (p.H54R), *MATR3* (p.K594I), *ALS2* (p.T787R), and *TARDBP* (p.L248F)—for detailed analysis of regional and cell-type distributions. Amplicon sequencing across multiple CNS regions showed that *MATR3* (p.K594I) and *TARDBP* (p.L248F) were restricted to the primary motor cortex (Fig. 4d and Supplementary Table 8), whereas *TIA1* (p.H54R) and *ALS2* (p.T787R) exhibited their highest VAFs in the spinal cord (2.16% and 0.97%, respectively), where they were originally identified, and were detected at much lower levels in other brain regions (Fig. 4d and Supplementary Table 8). All four somatic SNVs were absent from the cerebellum. The ultra-low VAFs and focal distribution of these variants suggest that they probably arose late in development and were thus likely CNS-restricted. Together with the enrichment of exonic and protein-altering somatic variants in disease-affected tissue regions, these findings also support the focal onset of ALS at the genetic level in these sporadic cases. Cells carrying damaging somatic variants could form initial lesions, likely TDP-43 inclusions, in UMNs and LMNs, with pathology ultimately spreading to other regions of the motor system that lack or carry exceedingly low levels of the variant, but nonetheless show robust post-mortem pathology otherwise indistinguishable from germline cases. Consistent with this hypothesis, the case harboring the *TARDBP* (p.L248F) variant, showed the highest phospho-TDP-43 (pTDP-43) level in the primary motor cortex, where the variant was detected (Extended Data Fig. 7). Quantification across seven brain regions revealed significant regional differences in pTDP-43 levels (*F* = 3.20, *P* = 0.0099, one-way ANOVA), with the motor cortex exhibiting significantly higher pathology compared to the hippocampus (*P* = 0.0042), middle temporal gyrus (*P* = 0.040), and occipital cortex (*P* = 0.032) in post hoc Tukey’s HSD tests. These results support the idea that focal somatic variants may initiate pathology that later spreads to broader brain regions.

We then assessed the distribution of these four somatic SNVs across cell types by performing amplicon sequencing of DNA from neuronal (NeuN+), glial (NeuN-), diploid, polyploid, and hypodiploid nuclei isolated by fluorescence-activated nuclei sorting (FANS) from the regions in which the variants were originally identified (Extended Data Fig. 8a,b). Interestingly, *TIA1* (p.H54R), *MATR3* (p.K594I), and *ALS2* (p.T787R) were enriched in hypodiploid nuclei (Extended Data Fig. 8c), which likely represent apoptotic cells (Supplementary Note)^49–52^, suggesting a potential association with cell death. Surprisingly, these three variants were identified in all cell fractions, but were more enriched in non-neuronal than neuronal populations (Extended Data Fig. 8c,d). This finding could imply that neurons may exhibit a cell-type-specific vulnerability to damaging somatic variants in ALS/FTD genes. However, further research is needed to confirm and better understand these potential associations and mechanisms. In contrast, the *TARDBP* (p.L248F) variant was validated only in the original primary motor cortex sample at very low VAF (∼0.5%) by both amplicon sequencing and ddPCR, but was not detected in sorted cell fractions from an additional adjacent primary motor cortex sample, indicating a highly focal event.

### RNA-MosaicHunter identifies additional predicted deleterious somatic variants in bulk RNA-seq data of sALS cases

To complement our targeted sequencing of neurodegenerative genes, which identified predicted deleterious somatic variants in a small proportion of sALS and sFTD cases, we performed a transcriptome-wide screen for somatic variants using bulk RNA-seq data to explore whether previously unknown genes might cause disease in a mosaic state. We profiled predicted deleterious somatic variants across all expressed genes in bulk RNA-seq data from 789 postmortem brain and spinal cord tissue samples of 143 sALS cases and 23 age-matched controls generated by the New York Genome Center ALS Consortium (Supplementary Table 9; 81 sALS and 11 control cases overlapped with our MIP cohort) using RNA-MosaicHunter^53^, which detects clonal somatic variants from bulk RNA-seq data using a Bayesian probabilistic model. Due to limited bulk RNA-seq depth, this approach is sensitive only to somatic variants with VAFs > ∼5% and does not detect somatic variants at ultra-low levels.

Although we did not observe a significant increase in the burden of total or predicted deleterious somatic variants in germline-free sALS cases (Supplementary Fig. 3), we identified predicted deleterious somatic SNVs in *DYNC1H1* and *LMNA* in multiple CNS regions of two sALS cases lacking other pathogenic or predicted deleterious germline/somatic variants (Fig. 5a and Supplementary Table 10); both cases overlapped with our MIP cohort. Heterozygously acting, generally *de novo*, variants in *DYNC1H1* and *LMNA* have been found in patients with phenotypes resembling spinal muscular atrophy (SMA)^54–57^, a motor neuron disease genetically distinct but sharing some pathological overlap with ALS^58^. Both individuals presented with leg-onset ALS and predominant spinal cord TDP-43 pathology (Fig. 5a-c). Amplicon sequencing revealed broad CNS distribution of the *LMNA* (p.H566Y) variant (VAF 5.3-12.3%) and the *DYNC1H1* (p.R1962C) variant (VAF 0.1-5.2%), with the *DYNC1H1* variant extremely low in the cerebellum (0.1%), thoracic spinal cord (0.8%) and lumbar spinal cord (0.8%) (Fig. 5d and Supplementary Table 8). Notably, the *DYNC1H1* (p.R1962C) variant was undetectable in cultured fibroblasts from the patient (Supplementary Table 8), indicating a late developmental,

**Fig. 5.**
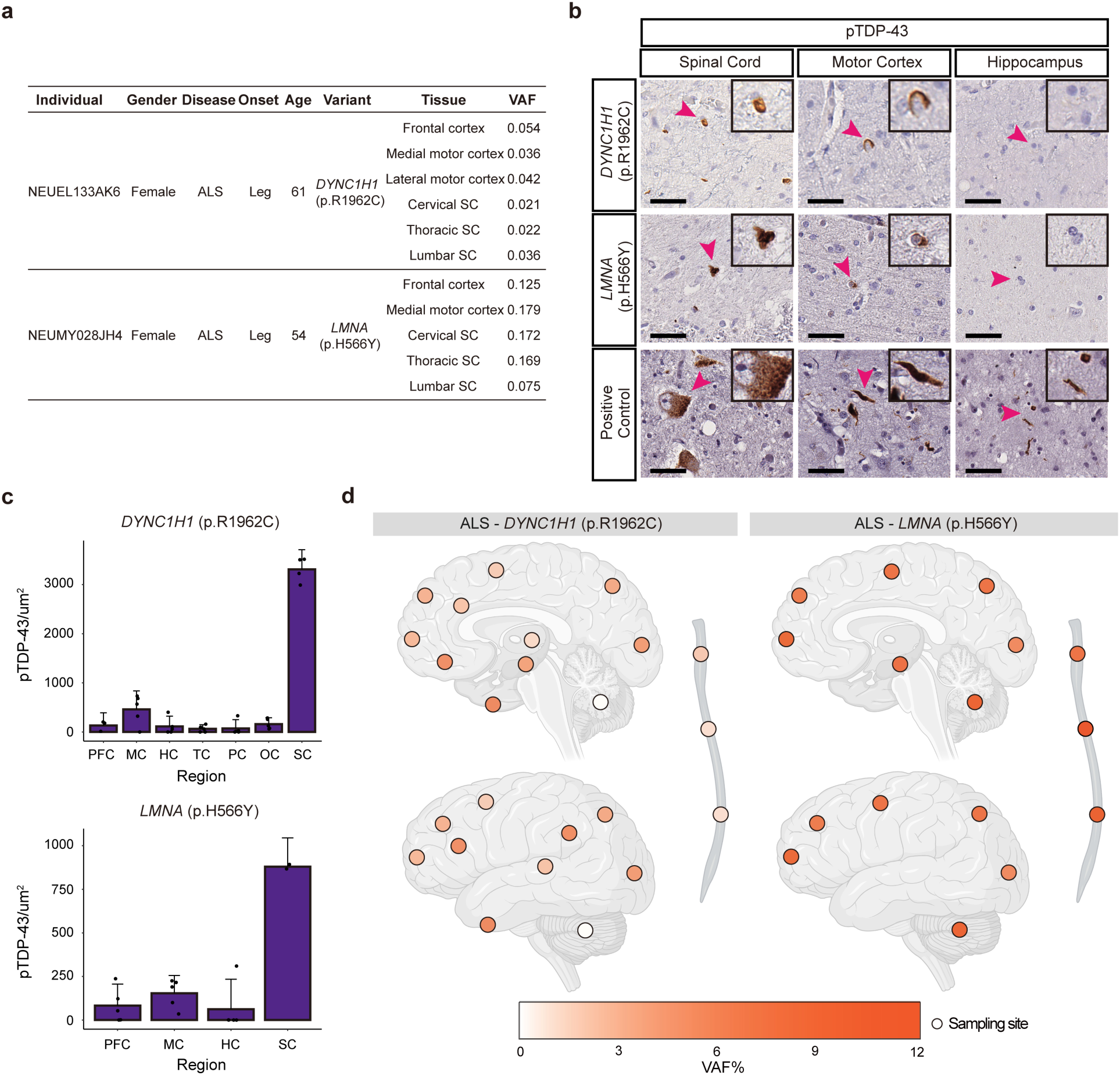
Somatic variants in *DYNC1H1* and *LMNA* in sALS. **a**, Two deleterious somatic SNVs that were shared by multiple tissue regions of the ALS cases. **b**, Sections of the lumbar spinal cord, motor cortex, and hippocampus of the two sALS cases stained with a phospho-TDP43 antibody. Scale bar, 40 μm. Arrowheads indicate the cells shown in the insets, which are magnified to twice their original size. **c**, Quantification of phospho-TDP43 staining of CNS tissue sections of the two sALS cases with *DYNC1H1* and *LMNA* somatic variants. For *DYNC1H1*, tissue-section-level biological replicates included 3 from PFC, 4 from SC, and 5 from MC, HC, TC, PC, and OC. For *LMNA*, tissue-section-level biological replicates included 2 from SC and 5 from PFC, MC, and HC. Bar graph, mean ± 95% CI. PFC, prefrontal cortex; MC, primary motor cortex; HC, hippocampus; TC, temporal cortex; PC, parietal cortex; OC, occipital cortex; SC, spinal cord. **d**, Regional distribution of VAFs of somatic variants in individual brains and spinal cords. Brain cortex is annotated by Brodmann areas. The color spectrum indicates the VAFs of somatic variants in amplicon sequencing. White indicates regions without the somatic variants. Created with BioRender.com.

CNS-restricted origin. The broad CNS distribution of these variants aligns with our previous observations that somatic variants above ∼5% VAFs are typically detected across the CNS^59^, with lower regional VAFs potentially reflecting selective neuronal loss. The *DYNC1H1* p.R1962C variant is highly pathogenic, as it completely abolishes dynein motor function *in vitro*^60^, and causes severe malformations of cortical development and delayed psychomotor development in patients carrying this germline variant^61,62^. Although the *LMNA* (p.H566Y) variant was not previously reported, germline *LMNA* variants cause autosomal dominant laminopathies with early lethality, including Hutchinson-Gilford progeria and congenital muscular dystrophy^63,64^. Thus, germline variants in both genes would ordinarily preclude ALS, but the mosaic state may allow for a normal early life followed by late-onset neurodegeneration. These findings suggest that further genome-wide exploration of brain tissue for somatic variants could reveal additional ALS genes that cause early lethality in the germline state.

### Somatic *C9orf72* repeat expansions detected by targeted long-read sequencing

The germline *C9orf72* repeat expansion is the most common genetic cause of ALS and FTD^27,28^. Repeat lengths > 30 are generally considered pathogenic, with most affected individuals carrying hundreds to thousands of repeats. Intermediate expansions with 20-30 repeats, confer an increased disease risk, while expansions with < 20 repeats are regarded as normal^27,65^. While, instability of pathogenic repeat expansions (> 30 repeats) across tissues within the same individual has been reported^66–68^, somatic expansion from non-expanded alleles has not been described. In our ALS and FTD cohort, the repeat-primed PCR analysis identified four ALS and FTD cases with both wild-type and expanded alleles showing reduced repeat peak heights^69^ (Extended Data Fig. 9), suggestive of somatic expansion. Targeted long-read sequencing in brain tissues from these cases confirmed the presence of both wild-type and expanded alleles with dozens to thousands of repeats. Remarkably, one FTD case carried two short wild-type alleles (4 and 9 repeats), together with highly expanded pathogenic alleles (Fig. 6a). Haplotype phasing based on heterozygous single-nucleotide polymorphisms (SNPs) flanking the repeat expansions showed that all expanded alleles shared the same SNP haplotype as the 4-repeat allele (Fig. 6b), strongly suggesting spontaneous somatic *C9orf72* repeat expansion of a non-expanded allele, though future analysis of more cases is needed to further characterize this possibility. In the remaining three cases, one wild-type allele and a spectrum of expanded alleles were detected, with minimum repeat sizes of 20, 56, and 64 repeats (Extended Data Fig. 10), consistent with somatic expansion from intermediate or short pathogenic alleles, although the possibility of contraction cannot be excluded. Together, these findings suggest that somatic *C9orf72* repeat expansions may contribute to sporadic ALS and FTD. Nevertheless, even long-read sequencing lacked sufficient sensitivity to comprehensively assess all germline-free cases (see Discussion), and the prevalence of somatic *C9orf72* expansions remains to be determined.

**Fig. 6.**
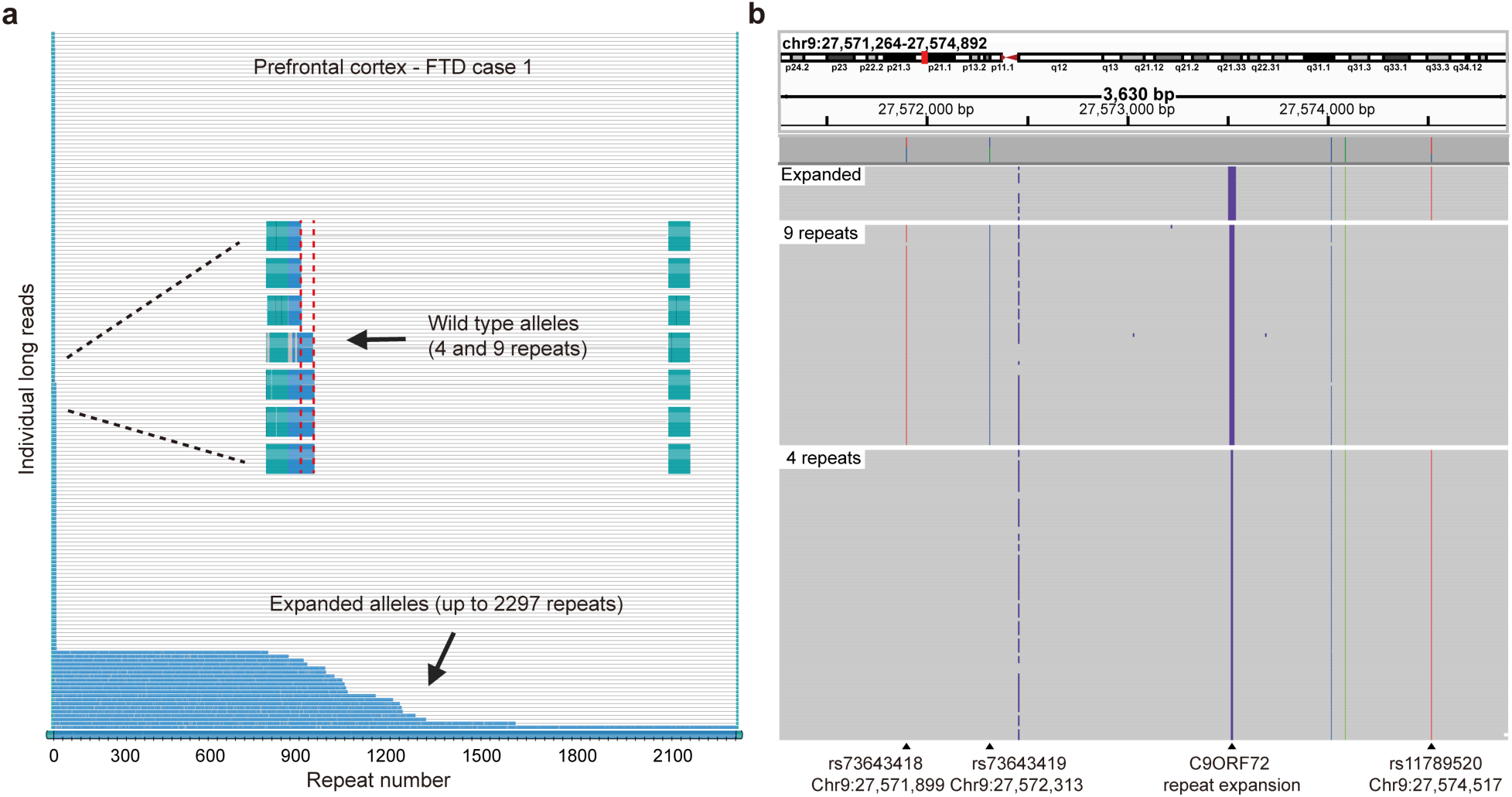
Somatic *C9orf72* repeat expansion in an FTD case. **a**, Waterfall plot showing targeted long-read sequencing results from the prefrontal cortex tissue of a sporadic FTD case. Two wild-type alleles with 4 and 9 repeats, respectively, are observed, along with somatic *C9orf72* repeat expansions ranging from 748 to 2,297 repeats. Each row represents an individual long read; the *C9orf72* repeat region is shown in blue, and the flanking regions are shown in green. Red dashed lines mark the sizes of the wild-type alleles. The *x*-axis denotes the number of GGGGCC hexanucleotide repeats. **b**, IGV screenshot showing haplotype phasing based on SNPs flanking the *C9orf72* repeat expansion. Long reads are grouped by repeat size category (expanded, 9 repeats, 4 repeats). SNPs rs73643418, rs73643419, and rs11789520 are heterozygous in this individual. All expanded alleles share the same SNP haplotype as the 4-repeat allele.

## Discussion

Our data provide several important insights into sALS and sFTD. First, approximately 30% of disease cases carried pathogenic or predicted deleterious germline variants in ALS/FTD genes, although our pathogenicity prediction might be overestimated. This advocates for a shift from family history-based to genetic testing-based disease classification, consistent with recently published guidelines for ALS genetic testing and counseling^70^. Second, a small but significant fraction (∼2.1%) of germline-free sporadic cases harbored predicted deleterious somatic variants in known ALS or FTD genes, with disease- and region-specific distributions in ALS, providing proof of concept for a contribution of somatic variants to disease pathogenesis. In addition, we identified genes associated with severe pediatric degenerative diseases that may contribute to ALS in the somatic state, broadening the spectrum of potential disease genes. Finally, targeted long-read sequencing revealed somatic *C9orf72* repeat expansions arising from wild-type, intermediate, and short pathogenic alleles in ALS and FTD cases.

While the case-control enrichment of somatic variants supports a pathogenic role, these variants occurred at surprisingly low VAFs and showed topographic restriction consistent with focal disease onset. Such variants likely arose late in development and were not shared by other tissue regions, exemplified by the *TARDBP* (p. L248F) variant detectable only at its site of discovery. The focal nature of these events would preclude detection by routine genetic testing with blood or other peripheral samples and support a mechanism by which degeneration may spread from a site containing mutant cells to anatomically connected regions, perhaps via intercellular transmission of TDP-43 proteinopathy^12–18,71,72^. The identification of predicted deleterious somatic variants in the primary motor cortex or spinal cord from individuals with ALS suggests that ALS may initiate in UMNs or LMNs before involving both.

While performed on a limited number of variants, our cell-type analysis revealed that several predicted deleterious somatic variants were shared across cell types and enriched in glia compared to neurons. However, this apparent glial enrichment may reflect preferential loss of neurons carrying these variants, consistent with their enrichment in hypodiploid cells, a population likely associated with apoptotic cell death. Together, these findings support a model in which deleterious somatic variants contribute to focal disease initiation, with subsequent neuronal loss leading to reduced VAFs over time.

Although only ∼2.1% of germline-free disease cases carried predicted deleterious somatic variants in ALS/FTD genes in our MIP sequencing data, this is likely greatly underestimated due to limited sensitivity for detecting ultra-low-VAF variants (Extended Data Fig. 1). Detecting such variants remains technically challenging^46^, and broader sampling across CNS regions is also limited. However, future duplex sequencing approaches promise the ability to define the minimal VAFs capable of initiating disease. On the other hand, several of our statistical comparisons approached nominal significance, highlighting the need for larger cohorts and more sensitive approaches.

While MIP sequencing did not allow for detection of somatic *C9orf72* repeat expansions in our samples, targeted long-read sequencing identified ALS and FTD cases with somatic expansions of varying repeat sizes, including an FTD case carrying two wild-type and highly expanded alleles, consistent with *de novo* expansion. These findings suggest that somatic *C9orf72* repeat expansions may contribute to sporadic ALS and FTD and could exhibit cell-type-specific patterns of expansion, analogous to somatic *HTT* CAG repeat expansion in striatal projection neurons^73^. However, the current targeted long-read sequencing approach preferentially enriches shorter alleles and provides limited flanking sequence, precluding accurate VAF estimation and robust haplotype phasing. Future targeted long-read sequencing with extended flanking coverage will be required to determine the origin and prevalence of somatic *C9orf72* repeat expansions.

Finally, the identification of somatic SNVs in *DYNC1H1* and *LMNA* suggests that genes predisposing to ALS and FTD in the somatic state may extend beyond those identified in germline studies. Although certain alleles in these genes cause motor neuron degeneration in the form of SMA, other alleles—including *DYNC1H1* p. R1962C^61,62^—cause severe pediatric disease that would preclude late-life ALS in the germline but may be compatible with disease in the somatic state. Similarly, enrichment in germline-free FTD cases emerged only when all targeted neurodegenerative genes were considered, hinting at a wider array of FTD-related genes. Together, these findings highlight the potential of future genome-wide approaches to uncover additional somatic genetic mechanisms and illuminate the topographic spread of pathology from focal origins.

## Acknowledgements

We thank the Massachusetts Alzheimer’s Disease Research Center, Oxford Brain Bank, Target ALS Foundation (Biobank Core Facility at St. Joseph’s Hospital and Barrow Neurological Institute, Georgetown Brain Bank, Eleanor and Lou Gehrig ALS Center at Columbia University and UCSD ALS bank) and NIH NeuroBioBank (Harvard Brain Tissue Resource Center, Mount Sinai/JJ Peters VA Medical Center NIH Brain and Tissue Repository, Brain Endowment Bank of University of Miami, University of Pittsburgh Neuropathology Brain Bank, University of Maryland Brian and Tissue Bank and UCLA Human Brain and Spinal Fluid Resource Center) for providing fresh frozen human tissues. We thank the Target ALS Human Postmortem Tissue Core, New York Genome Center for Genomics of Neurodegenerative Disease, Amyotrophic Lateral Sclerosis Association and TOW Foundation for providing the bulk RNA-seq data. We thank the donors and families for their contributions, and J. E. Neil and J. Gonzalez for assistance with tissue procurement. We thank the Research Computing group at Harvard Medical School and Boston Children’s Hospital. The brains in Figures 4 and 5 were illustrated by A. Lai with input from the authors. This work was supported by the PRMRP Discovery Award W81XWH2010028 (Z.Z.); the Edward R. and Anne G. Lefler Center postdoctoral fellowship (Z.Z.); the American Heart Association Career Development Award 23CDA1046074 (Z.Z.); the National Research Foundation of Korea (NRF) 2022R1C1C1010430 (Junho K.); RS-2023-00217881 (Junho K.); RS-2025-02215360 (Junho K.); R01 AG088082 (A.Y.H.); the Alzheimer’s Association research fellowship (A.Y.H.); R56 AG079857 (A.Y.H., C.A.W. and E.A.L.); a Cullen Education and Research Foundation Young Investigator Award from the Healey Center (M.N.); a Holloway Postdoctoral Fellowship from the Association for Frontotemporal Degeneration (M.N.); K08 AG065502 (M.B.M.); donors of the Alzheimer’s Disease Research program of the BrightFocus Foundation A20201292F (M.B.M.); the Doris Duke Charitable Foundation Clinical Scientist Development Award 2021183 (M.B.M.); K01 AG051791 (E.A.L.); the Suh Kyungbae Foundation (E.A.L.), DP2 AG072437 (E.A.L.); R01 NS032457 (C.A.W.); R01 AG070921 (C.A.W. and E.A.L.); a Massachusetts Alzheimer’s Disease Research Center pilot grant (C.L.-T. and C.A.W.); and the Allen Discovery Center program, a Paul G. Allen Frontiers Group advised program of the Paul G. Allen Family Foundation (C.A.W. and E.A.L.). C.L.-T. is supported by the Araminta Broch-Healey Endowed Chair in ALS. C.A.W. is an Investigator of the Howard Hughes Medical Institute. The funders had no role in the study design, data collection and analysis, decision to publish or preparation of the manuscript.

## Author Contributions

Z.Z., Junho K., A.Y.H., E.A.L., C.L.-T. and C.A.W. conceived and designed the study. Z.Z. performed tissue processing, MIP panel sequencing, cell sorting, amplicon sequencing, and targeted long-read sequencing. Junho K. performed bioinformatic analysis for MIP and validation data with assistance from R.D., T.S., and B.Z. A.Y.H. performed bioinformatic analysis for bulk RNA-seq data with assistance from J.P. and M.B. Jinhyeong K. performed bioinformatic analysis for long-read sequencing data. M.N. optimized and performed immunofluorescent imaging and quantification, and generated data shown in this manuscript. Z.Z., M.B.M. and R.D. designed the MIP panel. B.C., K.M., R.C.Y., and C.-Z.L. helped with tissue processing and amplicon sequencing. C.K. provided technical support for MIP sequencing. J.E.N. contributed tissue procurement and ethics expertise. T.O. and J.R. provided immunofluorescent images and interpretation of disease pathology. L.W.O. and O.A. provided fresh frozen human tissues and interpretation of disease pathology. C.L.-T., E.A.L. and C.A.W. supervised the study. Z.Z., Junho K., A.Y.H., C.L.-T., E.A.L. and C.A.W., wrote the manuscript.

## Methods

### Tissue sources and sample preparation

Fresh frozen postmortem human brain and spinal cord tissues were collected by the Massachusetts Alzheimer’s Disease Research Center, Oxford Brain Bank, Target ALS Foundation, and NIH NeuroBioBank (Supplementary Table 1) according to their respective institutional protocols, written authorization and informed consent; they were subsequently obtained for this study with the approval of the Boston Children’s Hospital Institutional Review Board. Research on these deidentified specimens and data was performed at Boston Children’s Hospital with approval from the Committee on Clinical Investigation. Sporadic ALS and FTD cases were selected based on available clinical records. ALS and FTD cases without clear recording of family histories were also selected if the age of death was above 45 years old. gDNA of these tissue samples was extracted using the EZ1 Advanced XL (Qiagen) system followed by an additional purification using AMPure XP beads (Beckman Coulter).

### MIP panel design

A double-stranded DNA MIP panel targeting 1.4 Mb across exons and exon-intron junctions of 88 neurodegenerative genes was designed using custom scripts incorporating MIPgen^74^ using the human reference genome, hg19, with Mly1 restriction sites masked with ‘N’ using bedtools. The final panel of 26,439 MIPs captures an average fragment length of 209 bp, including the extension and ligation arms to ensure overlapping of the forward and reverse sequencing read. The panel successfully targets 92.7% of bases including flanking intronic regions, with > 98% of exonic bases covered with an average of at least 2 unique MIPs. All MIPs were designed to include a custom backbone consisting of primer binding sites and dual 5-nt unique molecular identifier (UMI). MIPs were rebalanced in the pool based on the percent of GC content within the regions to improve coverage at GC-rich regions. We increased the copied MIPs that bind to GC-rich regions with the following criteria: 60-70% GC = 2 copies; 70-80% GC = 5 copies; 80-90% GC = 8 copies; > 90% GC = 10 copies. Common primer binding and Mly1 restriction enzyme sites were added to both ends of the oligo sequences to enable blunt-end removal of the primer binding sites. The forward and reverse compliment sequences were printed into a single ssDNA pool by CustomArray (Bothell, WA). The resulting panel was amplified at a low cycle number (12×), digested with Mly1 enzyme for 12 h at 37 °C, and purified using Qiagen Nucleotide removal kit.

### MIP capture and library construction

Two hundred fifty ng of gDNA was first hybridized in a 15 μl reaction with 1.5 μl of Ampligase® 10× Reaction Buffer (VWR), 1.5 μl of the reverse blocking oligo (5’-NNNNGAAGTCGAAGGGCTATAGGCTGCCATCACANNNN-3’) and the MIP pool at 63 nM for 10 min at 95 °C and 24 h at 60 °C. Gap-fill/ligation was then performed by adding 1 unit of Phusion™ High-Fidelity DNA Polymerase (Thermo Fisher), 4 units of Ampligase® DNA Ligase (Epicentre), 0.2 μl of Ampligase® 10× Reaction Buffer, 0.6 μl of dNTPs (10 mM) and 1 μl of nuclease-free water to the MIP capture product and incubated at 60 °C for 1 h. For exonuclease digestion, 50 units of Exonuclease III (Thermo Fisher), 10 units of Exonuclease I (Thermo Fisher), 0.2 μl of Ampligase® 10× Reaction Buffer (VWR), and 2.05 μl of nuclease-free water was added to the Gap-fill/ligation product, which was incubated for 40 min at 37 °C and 5 min at 95 °C. Ten μl of the captured library is amplified in a 50 μl final reaction by adding 1 unit of Phusion Hot Start II DNA Polymerase (Thermo Fisher), 10 μl of 5× HF buffer, 1 μl of dNTPs (10mM), 1 μl of the universal MIP barcode forward primer (10 μM), 1 μl of the individual barcode reserve primer (10 μM) and 26.5 μl of nuclease-free water. MIP library amplification was then performed under the following conditions: 98 °C for 30 s; 16 cycles of 98 °C for 10 s, 60 °C for 30 s and 72 °C for 30 s; 72 °C for 2 min. MIP library was then purified using 2× AMPure XP Beads (Beckman Coulter,) and quantified by the Quant-iT™ dsDNA Assay HS Kit (Thermo Fisher). Ninety-six MIP libraries were pooled together and sequenced on one lane of Illumina Hiseq X.

### Pre-processing and read mapping of MIP sequencing data

MIP sequencing primers were removed first from the raw FASTQ files using Cutadapt^75^ (v2.4; 5’ adapter of the first read: CATACGAGATCCGTAATCGGGAAGCTGAAG; 3’ adapter of the first read: ACACTACCGTCGGATCGTGCGTGT; 5’ adapter of the second read: GCTAAGGGCCTAACTGGCCGCTTCACTG; 3’ adapter of the second read: CTTCAGCTTCCCGATTACGGATCTCGTATG). Trimmed reads were aligned to the human reference genome (GRCh37) using BWA-mem^76^ (v0.7.15) and sorting and indexing were performed using samtools^77^ (v1.3.1). From the aligned BAM file, off-target reads were removed by checking the overlaps with the target regions using bedtools^78^ (v2.26.0). MIP arm regions were masked by soft-clipping for each read using BAMClipper^79^ (v1.0.1). UMI information was extracted, and then mapped reads were deduplicated based on the mapping coordinate and the shared UMI using UMI-tools^80^ (v1.0.0). Base quality score recalibration and local realignment were performed using the Genome Analysis Toolkit (GATK, v3.7)^81^, generating final analysis-ready BAMs.

### Variant calling for germline variants

Initial candidates of germline SNVs and indels were identified using GATK HaplotypeCaller with default parameter settings. Low-quality candidates were filtered out if any of the following conditions is not satisfied: (1) ≥ 10 variant-supporting reads; (2) ≥ 20 total read-depth at the variant site; (3) VAF ≥ 0.3; (4) GATK QUAL ≥ 50, and; (5) identified in all brain regions from the same individual except for the samples failed to cover the variant site (<10 reads). Possible pathogenic germline variants were further selected by satisfying all the following conditions: (1) present in less than 0.1% of the population in any ancestry group of public databases including dbSNP^82^, the 1000 Genomes Project^83^, the Exome Aggregation Consortium (ExAC)^84^, the Genome Aggregation Database (gnomAD)^85^, the NHLBI Exome Sequencing Project (ESP6500)^86^, the Greater Middle East variome project (GME)^87^, and Kaviar database^88^; (2) candidates observed only in disease or control groups but not in both; (3) possible protein-altering candidates (missense, nonsense, frame-shift, or splicing variants), and; (4) affecting 30 ALS- and FTD-related genes. Deleteriousness prediction module (see computational prediction of variant deleteriousness section below) was then applied to the remaining candidates, and predicted deleterious variants were reported as final deleterious germline variants. ANNOVAR^26^ was used to annotate the genomic region, affected genes, population allele frequency, and exonic variant functions. SpliceAI^89^ was additionally utilized to identify more splice-altering variants. Candidates with delta score > 0.5 were considered to be possible splicing variants.

### Repeat-primed PCR assay for *C9orf72* repeat expansion genotyping

Repeat-primed PCR (RP-PCR) of the *C9orf72* repeat expansion was performed in a 30 μl PCR reaction with 150 ng of gDNA, 15 μl of 2× FastStart™ PCR Master (Roche), 2 μl of DMSO, 5 μl of 5× Q-solution (Qiagen), 1 μl of 5 mM 7-deaza-dGTP (NEB), 1 μl of 25 mM MgCl_2_ (Qiagen) and 1 μl of the primer mix (40 μM of the Forward primer: 5’-/56-FAM/AGTCGCTAGAGGCGAAAGC-3’; 20 μM of the Reverse primer: 5’-TACGCATCCCAGTTTGAGACGGGGGCCGGGGCCGGGGCCGGGG-3’; 40 μM of the Anchor/tail primer: 5’-TACGCATCCCAGTTTGAGACG-3’). The reaction was performed with touchdown PCR cycling conditions consisting of 15 min at 95 °C, followed by cycles of 94 °C for 1 min, annealing starting at 70 °C for 1 min, and extension at 72 °C for 3 min, ending with a final extension step of 10 min at 72 °C. The annealing temperature was decreased in 2 °C steps as follows: 70 °C for two cycles, 68 °C for three cycles, 66 °C for four cycles, 64 °C for five cycles, 62 °C for six cycles, 60 °C for seven cycles, 58 °C for eight cycles, and 56 °C for five cycles. The RP-PCR products were separated by the SeqStudio Genetic Analyzer (Thermo Fisher) with the GeneScan™ 600 LIZ™ Dye Size Standard (Thermo Fisher). Results of fragment sizes were analyzed by Peak Scanner™ Software v1.0 (Thermo Fisher).

### Somatic variant calling from MIP sequencing data

Three different callers RePlow (v1.1.0)^38^, Mutect2 (v4.1.5)^39^, and Pisces (v5.2.11)^40^ were used to generate initial candidate sets. RePlow is optimized for detecting low-VAF somatic variants from deep targeted sequencing data. It generates a profile of background errors per substitution type and utilizes these distributions as priors in a Bayesian model to estimate the probability of a variant candidate being an error. MuTect2 is one of the most widely used somatic variant callers, particularly sensitive to detecting low-VAF variants in impure and heterogeneous samples. It employs a Bayesian classifier to evaluate the likelihood of a variant being genuine versus a sequencing error and applies multiple filters to reduce false positives. Pisces is specifically designed for detecting somatic mutations from amplicon sequencing data, particularly in cases where no matched control sample is available. It stitches paired-end reads into consensus reads and recalibrates variant quality scores specifically tuned to address amplification-related errors such as thermal damage or deamination, often observed in FFPE samples.

Each sample was analyzed by all three callers using single-sample mode. Default parameter settings were used except for the adjustments for disabling the coverage limit. Variants that passed all the filters from each caller were used to make three different initial sets. Candidates identified by only one caller were discarded, and those called at least two callers were retained as a double-call set. For indels, double-calls between Mutect2 and Pisces were used as somatic indel candidates since RePlow does not support indel detection. For SNVs, among double-calls Mutect2-Pisces pairs were additionally filtered out due to high false positive rates and low validation rates in the benchmarking data set (Extended Data Fig. 3d). Remaining RePlow-based SNV double-calls and indel candidates were subject to multi-step variant filters to further remove false positive candidates.

Unlike germline variant calling, somatic variant calling aims to reliably detect low-VAF variants up to ∼0.5%, which requires enough supporting evidence to control the false positive rate. Calling thresholds such as variant-supporting read count, read-depth at the variant site, and average base-call quality were determined based on the benchmarking data. Somatic variants were selected satisfying all the following conditions: (1) ≥ 50 total read-depth at the variant site; (2) ≥ 15 variant-supporting reads excluding the reads with the variant allele on their probe-arm regions; (3) > 30 average base-call quality of variant allele; (4) ≥ 2 different types of variant-supporting amplicons; (5) 0.001 ≤ VAF ≤ 0.4; (6) ≤ 3 variant candidates within 20-bp window from the same sample; (7) present in less than 0.1% of the population in any ancestry group of public databases, and; (8) observed in no more than two different individuals.

Somatic variants were further annotated with similar criteria for selecting predicted deleterious germline variants. Among the final candidates, variants that are (1) observed only in disease or control groups but not in both, (2) possible protein-altering variants, and (3) affecting ALS- and FTD-related genes were selected and applied for the deleteriousness prediction module. ANNOVAR and SpliceAI were utilized to annotate variants with various genomic information and detect additional splice-altering variants, respectively.

### Computational prediction of variant deleteriousness

Deleteriousness prediction module was applied to filtered germline and somatic variants to refine the predicted deleterious (potentially pathogenic) candidate sets. Variants that were previously reported as benign/likely benign in the clinical databases (ClinVar^90^ and Human Gene Mutation Database^91^) were excluded from the predicted candidate set. Nonsense, frameshift, and canonical splicing variants (±1-2 splice sites) were assumed to disrupt gene function and were included in the predicted deleterious set. For missense variants, the dbNSFP database^92^ was utilized to adopt multiple computational algorithms (SIFT^93^, PolyPhen2^94^, LRT^95^, MutationTaster^96^, MutationAssessor^97^, FATHMM^98^, FATHMM-MKL^99^, PROVEAN^100^, MetaSVM^101^, MetaLR^101^), considering damaging effects at different levels such as biochemical property, protein structure, and evolutionary conservation. Categorical prediction results of each algorithm were delivered by ANNOVAR. A missense variant was selected to be predicted deleterious if at least three different algorithms predicted damaging effects (deleterious for SIFT, LRT, FATHMM, PROVEAN, MetaSVM and MetaLR; probably damaging for PolyPhen2; disease-causing for MutationTaster). Possibly/likely damaging predictions were excluded for more conservative selection. For ALS/FTD-related genes, previously reported inheritance patterns (dominant/recessive) were carefully checked. For recessive genes, two independent variants in the same gene were required to determine whether a given individual was affected by predicted deleterious variants.

### Benchmarking with spike-in datasets

Extracted gDNA from two Coriell cell lines (GM12878 and GM24695) were used to generate a spike-in data. Extracted DNA were mixed at five different levels to mimic low-level somatic variants, targeting the VAFs of 0.5%, 1%, 2.5%, 5%, and 10%. Genomic DNA from GM12878 cells was spiked into DNA from GM24695, therefore unique germline SNPs in GM12878 were served as somatic variants. Genomic position and genotype information for germline SNPs of Coriell samples were obtained from NIST high-confidence call sets^102^. A total of 165 SNPs (57 homozygous and 108 heterozygous SNPs) were initially covered by our designed MIP panel. Among them, we only utilized heterozygous SNPs for benchmarking to accurately reflect target VAFs. Additionally, we excluded the spike-in variants from benchmarking if (1) a given site could be covered by only one amplicon, or (2) have read-depth of less than 100, as these are low-quality variants unable to be detected due to inherent issues with panel design rather than the variant calling process. RePlow, Mutect2, Pisces, and their combinations were tested. Detected variants not in the benchmark set were considered to be false positives, except for GM24695 germline SNPs and excluded spike-in variants.

### Somatic variant calling from bulk RNA-seq data

Raw bam files of bulk RNA-seq and matched WGS data for sALS and control cases of the New York Genome Center ALS Consortium were obtained from the New York Genome Center. RNA-seq reads extracted from raw bam files were aligned to the GRCh38 human reference genome by STAR (v2.5.0a)^103^ in the two-pass mode with the reference gene annotation (Gencode version 39). The aligned bam files were processed by Picard (v1.138) to remove duplicates, and then by GATK (v3.6)^104^ for SplitNCigarReads, indel realignment, and base quality recalibration. We further excluded reads that were improperly paired or with ambiguous alignment.

Somatic SNVs were called by RNA-MosaicHunter (v1.0) with default parameters. Derived from MosaicHunter^105^, which was designed for somatic variant calling in DNA sequencing, RNA-MosaicHunter incorporates a Bayesian genotyper and a series of empirical filters to systematically distinguish somatic variants from technical artifacts and germline variants, with 59% sensitivity and 94% precision benchmarked using cancer datasets. Specifically, germline variants identified from the matched WGS data from the same individual were excluded. We excluded A-to-G candidates because they are most likely led by the widespread A-to-I(G) RNA editing events in the human genome. To remove recurrent artifacts, we only considered exonic candidates that were called in one or two individuals. We further excluded candidates present in human polymorphism databases including dbSNP^82^, the 1000 Genomes Project^106^, the Exome Sequencing Project^107^, and the Exome Aggregation Consortium^108^.

### Long-read targeted sequencing

The PureTarget kit (PacBio) was used to prepare sequencing libraries with 2 µg of high-molecular-weight DNA following the manufacturer’s protocol. Pooled libraries were sequenced on one Revio SMRT Cell. Raw HiFi reads were aligned to the human reference genome (GRCh38) using minimap2 (v2.28)^109^, and sorting and indexing were performed using samtools (v1.3.1)^77^. The Tandem Repeat Genotyping Tool (TRGT, v1.1.1)^110^ was used to determine the *C9orf72* repeat counts for each sample (chr9:27573528-27573546, (GGGGCC)n). A waterfall plot was generated using TRVZ to visualize repeat counts with mosaicism.

### Nuclei isolation and whole genome amplification

Isolation of total (DAPI+), neuronal (NeuN+), non-neuronal (NeuN-), and damaged (low DAPI) nuclei were achieved by FANS together with nuclear staining of NeuN (Millipore, MAB377, clone A60, 1:1,500) and DAPI following a previously published study^111^. Five hundred nuclei of each cell population were sorted into wells of 96-well plates.

Sorted nuclei were subjected to genome amplification using the Primary Template-directed Amplification kit (BioSkryb, 100136) following the manufacturer’s protocol.

### Amplicon sequencing

Primer sets targeting each identified somatic SNV were designed using BatchPrimer3 (Supplementary Table 11). Amplicon was amplified for 25 cycles in a 50 μl PCR reaction with 50 ng of gDNA, 1 unit of Phusion Hot Start II DNA Polymerase (Thermo Fisher), 10 μl of 5× HF buffer, 1 μl of dNTPs (10 mM) and 10 μl of each primer (10 μM). Amplicon PCR products were then purified by a 0.65× + 1.05× double size selection with AMPure XP Beads (Beckman Coulter, A63882). Purified amplicons were then pooled based on the concentrations measured by the Quant-iT™ dsDNA Assay HS Kit (Thermo Fisher) and sequenced using Amplicon-EZ (Genewiz).

### Immunohistochemistry

Immunohistochemistry was performed using DAB (3,3’-Diaminobenzidine) detection following published protocols^112^. Briefly, 7-µm formalin-fixed, paraffin-embedded (FFPE) sections were dewaxed using citrisolve, before being rehydrated through decreasing concentrations of ethanol. Antigen retrieval was performed using sodium citrate buffer pH 6.0 at 121 °C for 15 min. Endogenous peroxidases were blocked using 3% hydrogen peroxide solution, and non-specific binding was blocked using 10% normal goat serum. Sections were then incubated overnight at 4 °C with primary antibody (pTDP-43 mouse polyclonal, CosmoBio CAC-TIP-PTD-P03, 1:10,000). After washing with TBS-Triton, sections were incubated with a Horseradish peroxidase (HRP)-conjugated Goat anti-mouse secondary (Dako) for 1 h at room temperature. HRP signal was detected using DAB substrate (Dako) applied for 15 min. Counterstaining was performed using Coles hematoxylin for 1 min. Sections were then dehydrated, cleared using citrisolve, and mounted using glass coverslips. All sections were viewed using a Leica upright light microscope and assessed for section quality prior to whole-slide digital scanning.

### Quantification of p-TDP43 burden by immunohistochemistry

Stained sections were scanned using a NanoZoomer whole-slide digital imager at 40X magnification. Images were then visualized and quantified using QuPath image analysis software and algorithms following published protocols^112^. Briefly, for cortical/cerebellar sections 5 ROI measuring 3 mm^2^ (1000 × 3000 µm) were placed equidistantly around a single gyrus with the short end of the ROI placed at the pial surface. Pathology was then quantified using a positive pixel count within each ROI and measurements were averaged to provide an output of positive pixels/mm^2^. For spinal cord sections, square ROI (2.25 mm^2^) was placed on each side of the central canal within the anterior horn and measurements were averaged.

### Data Availability

The bulk RNA-seq data generated by the NYGC ALS Consortium are available through controlled access via the Target ALS Data Portal (https://dataengine.targetals.org/). Access requires acceptance of the Target ALS Data Use Agreement and submission of a data access request through the portal application process. Additional details are provided in the Target ALS Data Portal User Manual. The MIP-based targeted sequencing data and long-read sequencing data generated in this study have been deposited in dbGaP under accession number phs003530, with access governed by human subject privacy regulations. Germline and somatic variants identified and validated in this study are listed in the supplementary tables.

### Code Availability

The source code and default configuration file of RNA-MosaicHunter have been published and are available at https://github.com/AugustHuang/RNA-MosaicHunter. Other custom analysis scripts used in this study are publicly available at GitHub (https://github.com/kimjh607/ALS-FTD-somatic-mosaicism) and archived at Zenodo (https://doi.org/10.5281/zenodo.18682277).

## References

1. Ferrari, R., Kapogiannis, D., Huey, E. D. & Momeni, P. FTD and ALS: a tale of two diseases. Curr. Alzheimer Res. 8, 273–294 (2011).

2. Saxon, J. A. et al. Examining the language and behavioural profile in FTD and ALS-FTD. J. Neurol. Neurosurg. Psychiatry 88, 675–680 (2017).

3. Lagier-Tourenne, C., Polymenidou, M. & Cleveland, D. W. TDP-43 and FUS/TLS: emerging roles in RNA processing and neurodegeneration. Hum. Mol. Genet. 19, R46–R64 (2010).

4. Ling, S. C., Polymenidou, M. & Cleveland, D. W. Converging mechanisms in ALS and FTD: disrupted RNA and protein homeostasis. Neuron 79, 416–438 (2013).

5. Ravits, J. M. & La Spada, A. R. ALS motor phenotype heterogeneity, focality, and spread: deconstructing motor neuron degeneration. Neurology 73, 805–811 (2009).

6. Kanouchi, T., Ohkubo, T. & Yokota, T. Can regional spreading of amyotrophic lateral sclerosis motor symptoms be explained by prion-like propagation? J. Neurol. Neurosurg. Psychiatry 83, 739–745 (2012).

7. Eisen, A., Kim, S. & Pant, B. Amyotrophic lateral sclerosis (ALS): a phylogenetic disease of the corticomotoneuron? Muscle Nerve 15, 219–224 (1992).

8. Chou, S. M. & Norris, F. H. Amyotrophic lateral sclerosis: lower motor neuron disease spreading to upper motor neurons. Muscle Nerve 16, 864–869 (1993).

9. Gromicho, M. et al. Spreading in ALS: the relative impact of upper and lower motor neuron involvement. Ann. Clin. Transl. Neurol. 7, 1181–1192 (2020).

10. Brettschneider, J. et al. Stages of pTDP-43 pathology in amyotrophic lateral sclerosis. Ann. Neurol. 74, 20–38 (2013).

11. Brettschneider, J. et al. Sequential distribution of pTDP-43 pathology in behavioral variant frontotemporal dementia (bvFTD). Acta Neuropathol. 127, 423–439 (2014).

12. Polymenidou, M. & Cleveland, D. W. Biological spectrum of amyotrophic lateral sclerosis prions. Cold Spring Harb. Perspect. Med. 7, a024133 (2017).

13. Porta, S. et al. Patient-derived frontotemporal lobar degeneration brain extracts induce formation and spreading of TDP-43 pathology in vivo. Nat. Commun. 9, 4220 (2018).

14. Laferriere, F. et al. TDP-43 extracted from frontotemporal lobar degeneration subject brains displays distinct aggregate assemblies and neurotoxic effects reflecting disease progression rates. Nat. Neurosci. 22, 65–77 (2019).

15. Peng, C., Trojanowski, J. Q. & Lee, V. M. Protein transmission in neurodegenerative disease. Nat. Rev. Neurol. 16, 199–212 (2020).

16. De Rossi, P. et al. FTLD-TDP assemblies seed neoaggregates with subtype-specific features via a prion-like cascade. EMBO Rep. 22, e53877 (2021).

17. Tamaki, Y. et al. Spinal cord extracts of amyotrophic lateral sclerosis spread TDP-43 pathology in cerebral organoids. PLoS Genet. 19, e1010606 (2023).

18. Kumar, S. T. et al. Seeding the aggregation of TDP-43 requires post-fibrillization proteolytic cleavage. Nat. Neurosci. 26, 983–996 (2023).

19. Rosen, D. R. et al. Mutations in Cu/Zn superoxide dismutase gene are associated with familial amyotrophic lateral sclerosis. Nature 362, 59–62 (1993).

20. Turner, M. R. et al. Controversies and priorities in amyotrophic lateral sclerosis. Lancet Neurol. 12, 310–322 (2013).

21. Wang, H. et al. Smoking and risk of amyotrophic lateral sclerosis: a pooled analysis of 5 prospective cohorts. Arch. Neurol. 68, 207–213 (2011).

22. Armon, C. Acquired nucleic acid changes may trigger sporadic amyotrophic lateral sclerosis. Muscle Nerve 32, 373–377 (2005).

23. Jamuar, S. S. et al. Somatic mutations in cerebral cortical malformations. N. Engl. J. Med. 371, 733–743 (2014).

24. Proukakis, C. Somatic mutations in neurodegeneration: an update. Neurobiol. Dis. 144, 105021 (2020).

25. Hardenbol, P. et al. Multiplexed genotyping with sequence-tagged molecular inversion probes. Nat. Biotechnol. 21, 673–678 (2003).

26. Wang, K., Li, M. & Hakonarson, H. ANNOVAR: functional annotation of genetic variants from high-throughput sequencing data. Nucleic Acids Res. 38, e164 (2010).

27. Renton, A. E. et al. A hexanucleotide repeat expansion in *C9ORF72* is the cause of chromosome 9p21-linked ALS-FTD. Neuron 72, 257–268 (2011).

28. DeJesus-Hernandez, M. et al. Expanded GGGGCC hexanucleotide repeat in noncoding region of *C9ORF72* causes chromosome 9p-linked FTD and ALS. Neuron 72, 245–256 (2011).

29. Majounie, E. et al. Frequency of the C9orf72 hexanucleotide repeat expansion in patients with amyotrophic lateral sclerosis and frontotemporal dementia: a cross-sectional study. Lancet Neurol. 11, 323–330 (2012).

30. Byrne, S. et al. Cognitive and clinical characteristics of patients with amyotrophic lateral sclerosis carrying a C9orf72 repeat expansion: a population-based cohort study. Lancet Neurol. 11, 232–240 (2012).

31. Mahoney, C. J. et al. Frontotemporal dementia with the C9ORF72 hexanucleotide repeat expansion: clinical, neuroanatomical and neuropathological features. Brain 135, 736–750 (2012).

32. Baker, M. et al. Mutations in progranulin cause tau-negative frontotemporal dementia linked to chromosome 17. Nature 442, 916–919 (2006).

33. Cruts, M. et al. Null mutations in progranulin cause ubiquitin-positive frontotemporal dementia linked to chromosome 17q21. Nature 442, 920–924 (2006).

34. van Blitterswijk, M. et al. Evidence for an oligogenic basis of amyotrophic lateral sclerosis. Hum. Mol. Genet. 21, 3776–3784 (2012).

35. Testi, S., Tamburin, S., Zanette, G. & Fabrizi, G. M. Co-occurrence of the *C9ORF72* expansion and a novel GRN mutation in a family with alternative expression of frontotemporal dementia and amyotrophic lateral sclerosis. J. Alzheimers Dis. 44, 49–56 (2015).

36. Kuuluvainen, L. et al. Oligogenic basis of sporadic ALS: the example of SOD1 p.Ala90Val mutation. Neurol. Genet. 5, e335 (2019).

37. Goutman, S. A. et al. Emerging insights into the complex genetics and pathophysiology of amyotrophic lateral sclerosis. Lancet Neurol. 21, 465–479 (2022).

38. Kim, J. et al. The use of technical replication for detection of low-level somatic mutations in next-generation sequencing. Nat. Commun. 10, 1047 (2019).

39. Benjamin, D. et al. Calling somatic SNVs and indels with Mutect2. bioRxiv, 861054 (2019).

40. Dunn, T. et al. Pisces: an accurate and versatile variant caller for somatic and germline next-generation sequencing data. Bioinformatics 35, 1579–1581 (2019).

41. Barnell, E. K. et al. Open-sourced CIViC annotation pipeline to identify and annotate clinically relevant variants using single-molecule molecular inversion probes. JCO Clin Cancer Inform 3, 1–12 (2019).

42. Biezuner, T., et al. An improved molecular inversion probe based targeted sequencing approach for low variant allele frequency. NAR Genom. Bioinform. 4, lqab125 (2022).

43. Bizzotto, S. et al. Landmarks of human embryonic development inscribed in somatic mutations. Science 371, 1249–1253 (2021).

44. Lee, J. et al. Mutalisk: a web-based somatic MUTation AnaLyIS toolKit for genomic, transcriptional and epigenomic signatures. Nucleic Acids Res. 46, W102–W108 (2018).

45. Chung, C. et al. Comprehensive multi-omic profiling of somatic mutations in malformations of cortical development. Nat. Genet. 55, 209–220 (2023).

46. Huang, A. Y. & Lee, E. A. Identification of somatic mutations from bulk and single-cell sequencing data. *Front*. Aging 2, 800380 (2021).

47. Hadano, S. et al. A gene encoding a putative GTPase regulator is mutated in familial amyotrophic lateral sclerosis 2. Nat. Genet. 29, 166–173 (2001).

48. Yang, Y. et al. The gene encoding alsin, a protein with three guanine-nucleotide exchange factor domains, is mutated in a form of recessive amyotrophic lateral sclerosis. Nat. Genet. 29, 160–165 (2001).

49. Darzynkiewicz, Z. et al. Features of apoptotic cells measured by flow cytometry. Cytometry 13, 795–808 (1992).

50. Arends, M. J., Morris, R. G. & Wyllie, A. H. Apoptosis. The role of the endonuclease. Am. J. Pathol. 136, 593–608 (1990).

51. Telford, W. G., King, L. E. & Fraker, P. J. Comparative evaluation of several DNA binding dyes in the detection of apoptosis-associated chromatin degradation by flow cytometry. Cytometry 13, 137–143 (1992).

52. Del Bino, G., Lassota, P. & Darzynkiewicz, Z. The S-phase cytotoxicity of camptothecin. Exp. Cell Res. 193, 27–35 (1991).

53. Huang, A. Y. et al. Accurate detection of somatic single-nucleotide variants from bulk RNA-seq data using RNA-MosaicHunter. Nucleic Acids Res. 54, gkaf1450 (2026).

54. Rudnik-Schoneborn, S. et al. Mutations of the LMNA gene can mimic autosomal dominant proximal spinal muscular atrophy. Neurogenetics 8, 137–142 (2007).

55. Harms, M.B. et al. Mutations in the tail domain of *DYNC1H1* cause dominant spinal muscular atrophy. Neurology 78, 1714–1720 (2012).

56. Tsurusaki, Y. et al. A *DYNC1H1* mutation causes a dominant spinal muscular atrophy with lower extremity predominance. Neurogenetics 13, 327–332 (2012).

57. Iwahara, N., Hisahara, S., Hayashi, T., Kawamata, J. & Shimohama, S. A novel lamin A/C gene mutation causing spinal muscular atrophy phenotype with cardiac involvement: report of one case. BMC Neurol. 15, 13 (2015).

58. Bowerman, M. et al. Pathogenic commonalities between spinal muscular atrophy and amyotrophic lateral sclerosis: Converging roads to therapeutic development. Eur. J. Med. Genet. 61, 685–698 (2018).

59. Lodato, M. A. et al. Somatic mutation in single human neurons tracks developmental and transcriptional history. Science 350, 94–98 (2015).

60. Hoang, H. T., Schlager, M. A., Carter, A. P. & Bullock, S. L. *DYNC1H1* mutations associated with neurological diseases compromise processivity of dynein-dynactin-cargo adaptor complexes. Proc. Natl. Acad. Sci. USA 114, E1597–E1606 (2017).

61. Poirier, K. et al. Mutations in *TUBG1*, *DYNC1H1*, *KIF5C* and *KIF2A* cause malformations of cortical development and microcephaly. Nat. Genet. 45, 639–647 (2013).

62. Yang, H. et al. De novo variants in the *DYNC1H1* gene associated with infantile spasms. Front. Neurol. 12, 733178 (2021).

63. Eriksson, M. et al. Recurrent de novo point mutations in lamin A cause Hutchinson-Gilford progeria syndrome. Nature 423, 293–298 (2003).

64. Quijano-Roy, S. et al. De novo *LMNA* mutations cause a new form of congenital muscular dystrophy. Ann. Neurol. 64, 177–186 (2008).

65. Breevoort, S., Gibson, S., Figueroa, K., Bromberg, M. & Pulst, S. Expanding clinical spectrum of *C9ORF72*-related disorders and promising therapeutic strategies: a Review. Neurol. Genet. 8, e670 (2022).

66. van Blitterswijk, M. et al. Association between repeat sizes and clinical and pathological characteristics in carriers of *C9ORF72* repeat expansions (Xpansize-72): a cross-sectional cohort study. Lancet Neurol. 12, 978–988 (2013).

67. DeJesus-Hernandez, M. et al. Long-read targeted sequencing uncovers clinicopathological associations for *C9orf72*-linked diseases. Brain 144, 1082–1088 (2021).

68. Udine, E. et al. Targeted long-read sequencing to quantify methylation of the *C9orf72* repeat expansion. Mol. Neurodegener. 19, 99 (2024).

69. Ross, J. P. et al. Somatic expansion of the *C9orf72* hexanucleotide repeat does not occur in ALS spinal cord tissues. Neurol. Genet. 5, e317 (2019).

70. Roggenbuck, J. et al. Evidence-based consensus guidelines for ALS genetic testing and counseling. Ann. Clin. Transl. Neurol. 10, 2074–2091 (2023).

71. Frank, S.A. Evolution in health and medicine Sackler colloquium: Somatic evolutionary genomics: mutations during development cause highly variable genetic mosaicism with risk of cancer and neurodegeneration. Proc. Natl. Acad. Sci. USA 107 Suppl 1, 1725–1730 (2010).

72. Pamphlett, R. The “somatic-spread” hypothesis for sporadic neurodegenerative diseases. Med. Hypotheses 77, 544–547 (2011).

73. Handsaker, R. E. et al. Long somatic DNA-repeat expansion drives neurodegeneration in Huntington’s disease. Cell 188, 623–639.e19 (2025).

## Methods-only References

74. Boyle, E. A., O’Roak, B. J., Martin, B. K., Kumar, A. & Shendure, J. MIPgen: optimized modeling and design of molecular inversion probes for targeted resequencing. Bioinformatics 30, 2670–2672 (2014).

75. Martin, M. Cutadapt removes adapter sequences from high-throughput sequencing reads. 2011 17, 3 (2011).

76. Li, H. Aligning sequence reads, clone sequences and assembly contigs with BWA-MEM. *arXiv preprint arXiv:1303*. 3997 (2013).

77. Danecek, P. et al. Twelve years of SAMtools and BCFtools. Gigascience 10, giab008 (2021).

78. Quinlan, A. R. & Hall, I. M. BEDTools: a flexible suite of utilities for comparing genomic features. Bioinformatics 26, 841–842 (2010).

79. Au, C. H., Ho, D. N., Kwong, A., Chan, T. L. & Ma, E. S. K. BAMClipper: removing primers from alignments to minimize false-negative mutations in amplicon next-generation sequencing. Sci. Rep. 7, 1567 (2017).

80. Smith, T., Heger, A. & Sudbery, I. UMI-tools: modeling sequencing errors in Unique Molecular Identifiers to improve quantification accuracy. Genome Res. 27, 491–499 (2017).

81. Van der Auwera, G. A. et al. From FastQ data to high confidence variant calls: the Genome Analysis Toolkit best practices pipeline. Curr. Protoc. Bioinformatics 43, 11.10.1–11.10.33 (2013).

82. Sherry, S. T. et al. dbSNP: the NCBI database of genetic variation. Nucleic Acids Res. 29, 308–11 (2001).

83. 1000 Genomes Project Consortium et al. A global reference for human genetic variation. Nature 526, 68–74 (2015).

84. Karczewski, K. J. et al. The ExAC browser: displaying reference data information from over 60 000 exomes. Nucleic Acids Res. 45, D840–D845 (2017).

85. Karczewski, K. J. et al. The mutational constraint spectrum quantified from variation in 141,456 humans. Nature 581, 434–443 (2020).

86. Fu, W. et al. Analysis of 6,515 exomes reveals the recent origin of most human protein-coding variants. Nature 493, 216–20 (2013).

87. Scott, E. M. et al. Characterization of Greater Middle Eastern genetic variation for enhanced disease gene discovery. Nat. Genet. 48, 1071–1076 (2016).

88. Glusman, G., Caballero, J., Mauldin, D. E., Hood, L. & Roach, J. C. Kaviar: an accessible system for testing SNV novelty. Bioinformatics 27, 3216–3217 (2011).

89. Jaganathan, K. et al. Predicting splicing from primary sequence with deep learning. Cell 176, 535–548.e24 (2019).

90. Landrum, M. J. et al. ClinVar: improving access to variant interpretations and supporting evidence. Nucleic Acids Res. 46, D1062–D1067 (2018).

91. Stenson, P. D. et al. Human Gene Mutation Database (HGMD): 2003 update. Hum. Mutat. 21, 577–581 (2003).

92. Liu, X., Wu, C., Li, C. & Boerwinkle, E. dbNSFP v3.0: a one-stop database of functional predictions and annotations for human nonsynonymous and splice-site SNVs. Hum. Mutat. 37, 235–41 (2016).

93. Kumar, P., Henikoff, S. & Ng, P. C. Predicting the effects of coding non-synonymous variants on protein function using the SIFT algorithm. Nat. Protoc. 4, 1073–1081 (2009).

94. Adzhubei, I. A. et al. A method and server for predicting damaging missense mutations. Nat Methods 7, 248–249 (2010).

95. Chun, S. & Fay, J. C. Identification of deleterious mutations within three human genomes. Genome Res. 19, 1553–1561 (2009).

96. Schwarz, J. M., Rodelsperger, C., Schuelke, M. & Seelow, D. MutationTaster evaluates disease-causing potential of sequence alterations. Nat. Methods 7, 575–576 (2010).

97. Reva, B., Antipin, Y. & Sander, C. Predicting the functional impact of protein mutations: application to cancer genomics. Nucleic Acids Res. 39, e118 (2011).

98. Shihab, H. A. et al. Predicting the functional, molecular, and phenotypic consequences of amino acid substitutions using hidden Markov models. Hum. Mutat. 34, 57–65 (2013).

99. Shihab, H. A. et al. An integrative approach to predicting the functional effects of non-coding and coding sequence variation. Bioinformatics 31, 1536–1543 (2015).

100. Choi, Y., Sims, G. E., Murphy, S., Miller, J. R. & Chan, A. P. Predicting the functional effect of amino acid substitutions and indels. PLoS One 7, e46688 (2012).

101. Dong, C. et al. Comparison and integration of deleteriousness prediction methods for nonsynonymous SNVs in whole exome sequencing studies. Hum. Mol. Genet. 24, 2125–2137 (2015).

102. Zook, J. M. et al. Integrating human sequence data sets provides a resource of benchmark SNP and indel genotype calls. Nat. Biotechnol. 32, 246–251 (2014).

103. Dobin, A. et al. STAR: ultrafast universal RNA-seq aligner. Bioinformatics 29, 15–21 (2013).

104. DePristo, M. A. et al. A framework for variation discovery and genotyping using next-generation DNA sequencing data. Nat. Genet. 43, 491–498 (2011).

105. Huang, A. Y. et al. MosaicHunter: accurate detection of postzygotic single-nucleotide mosaicism through next-generation sequencing of unpaired, trio, and paired samples. Nucleic Acids Res. 45, e76 (2017).

106. 1000 Genomes Project Consortium et al. An integrated map of genetic variation from 1,092 human genomes. Nature 491, 56–65 (2012).

107. Tennessen, J. A. et al. Evolution and functional impact of rare coding variation from deep sequencing of human exomes. Science 337, 64–69 (2012).

108. Lek, M. et al. Analysis of protein-coding genetic variation in 60,706 humans. Nature 536, 285–291 (2016).

109. Li, H. Minimap2: pairwise alignment for nucleotide sequences. Bioinformatics 34, 3094–3100 (2018).

110. Dolzhenko, E. et al. Characterization and visualization of tandem repeats at genome scale. Nat. Biotechnol. 42, 1606–1614 (2024).

111. Evrony, G. D. et al. Single-neuron sequencing analysis of L1 retrotransposition and somatic mutation in the human brain. Cell 151, 483–496 (2012).

112. Nolan, M. et al. Quantitative patterns of motor cortex proteinopathy across ALS genotypes. Acta Neuropathol Commun 8, 98 (2020).

