## Supplemental Information for "Somatic mosaicism in amyotrophic lateral sclerosis and frontotemporal dementia identifies focal mutations associated with widespread degeneration"

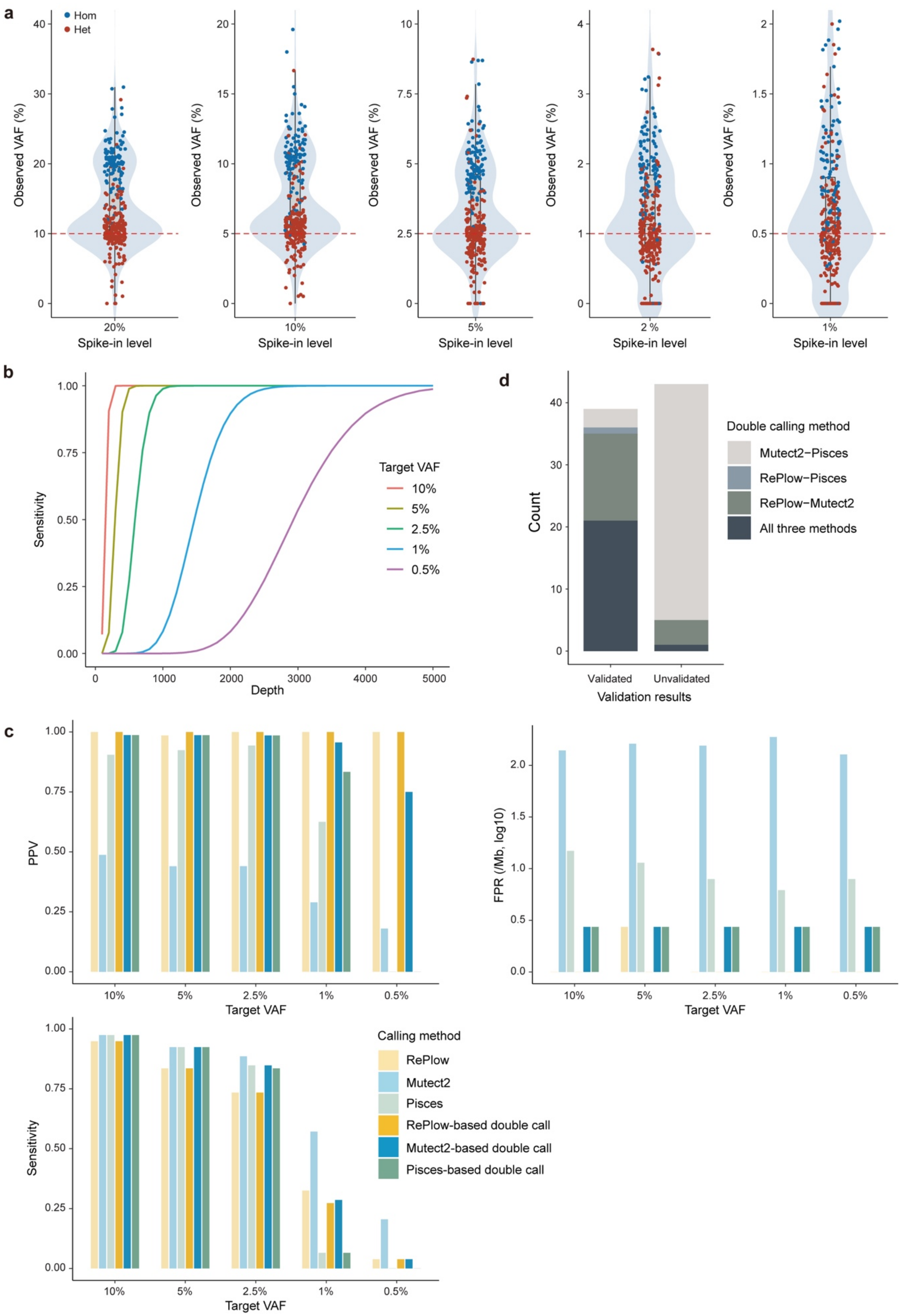

a
Hom
Het
Observed VAF (%)
Spike-in level
20%
10%
5%
2 %
1%
b
Sensitivity
Depth
Target VAF
10%
5%
2.5%
1%
0.5%
c
PPV
FPR (/Mb, log10)
Sensitivity
Target VAF
Calling method
RePlow
Mutect2
Pisces
RePlow-based double call
Mutect2-based double call
Pisces-based double call
d
Count
Validation results
Validated
Unvalidated
Double calling method
Mutect2−Pisces
RePlow−Pisces
RePlow−Mutect2
All three methods

**Extended Data Fig. 1: Evaluation of the variant calling method for somatic variant using spike-in and validation experiments.** (a) VAF distribution of 165 artificial variants from the MIP-sequenced spike-in data. Each violin plot denotes benchmark variants across spike-in mixtures at five different levels (20%, 10%, 5%, 2%, and 1%). Box plots indicate the median (center line) and interquartile range (25th–75th percentiles), with whiskers extending to the most extreme values within 1.5× the interquartile range. Each dot represents the VAF of an individual variant. Red dotted lines represent the target VAF of a given mixture. (b) Theoretical sensitivity of variant detection across varying VAF levels and sequencing depths under error-free conditions. (c) Benchmarking of three different callers and their combinations using spike-in data. Note that only heterozygous variants were utilized for benchmarking to specifically evaluate performance for the target VAFs. PPV, Positive predictive value; FPR, false positive rate. (d) Validation results of identified double-called variants using deep amplicon sequencing. Each color represents a different double-calling combination.

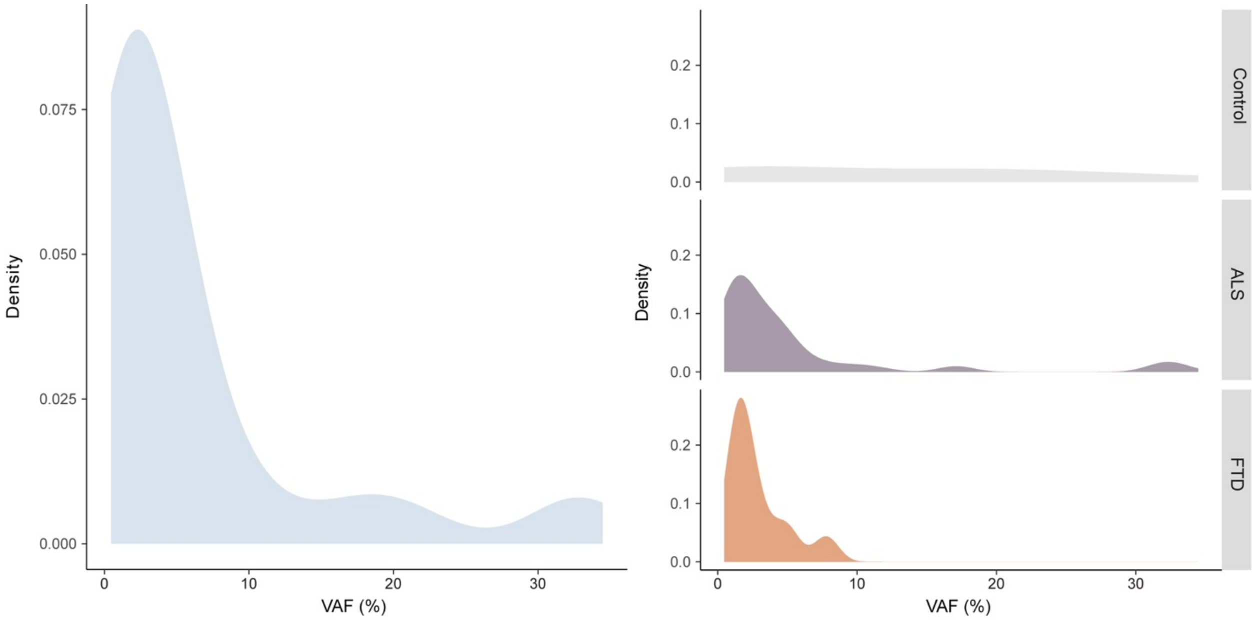


**Extended Data Fig. 2: Variant allele fraction (VAF) distribution of identified somatic variants.** VAF distribution of total somatic variants (left) and across different clinical conditions (right) are shown in density plot. All VAFs were used to draw the figure if a given variant was observed multiple times in different brain regions.

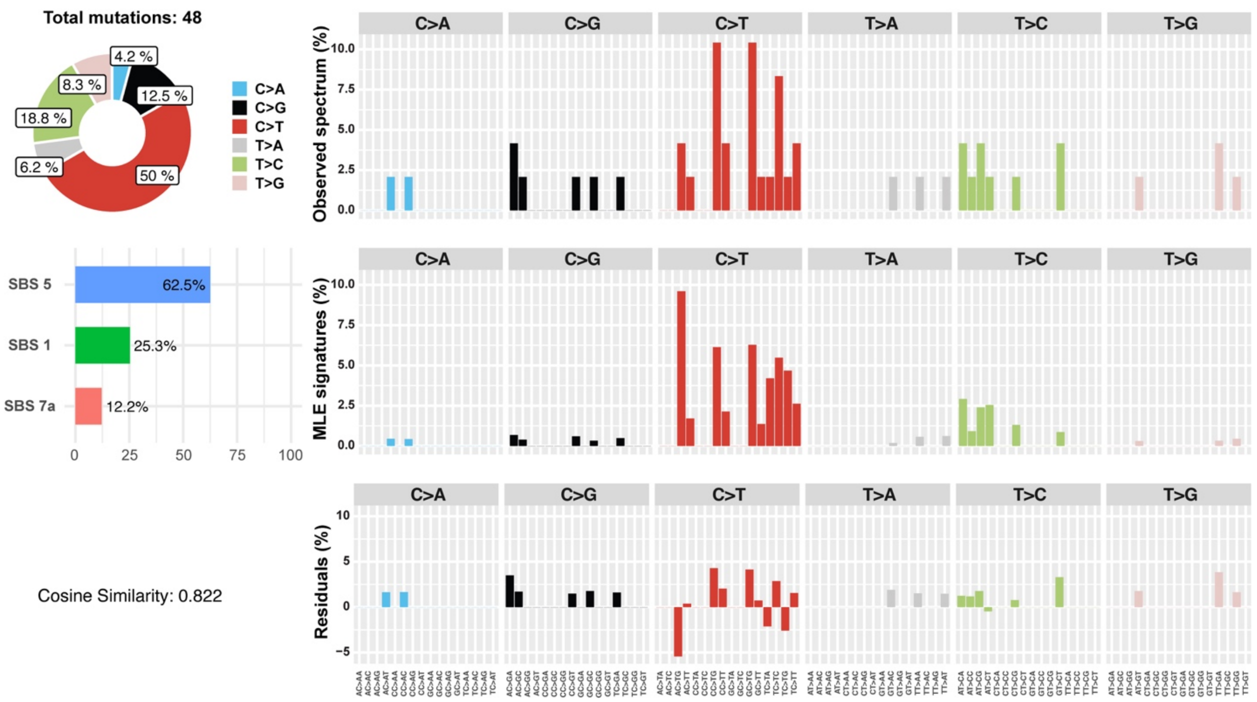


**Extended Data Fig. 3: Somatic SNV profile and underlying mutational processes.** Mutational signature analysis using Mutalisk revealed two clock-like signatures (SBS5 and SBS1) as major underlying mutational mechanisms. Note that Mutalisk accounts for SNVs only, resulting in a total of 48 unique variants obtained from germline-free individuals. SBS1 indicates mutations caused by deamination of methylated cytosine in cycling cells and is directly associated with stem cell division and mitosis.

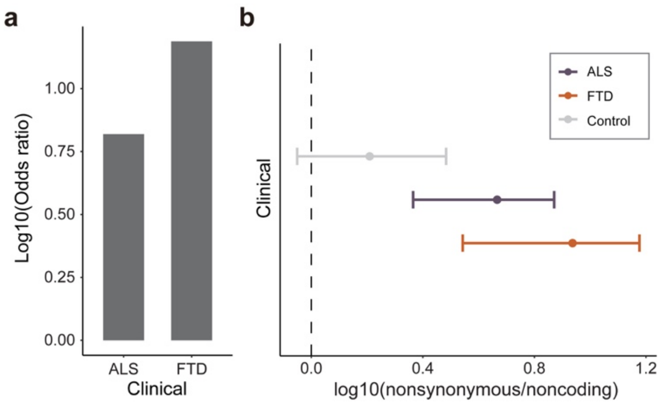


**Extended Data Fig. 4: Ratios of non-synonymous to non-coding variants in germline-free ALS and FTD cases.** (a) Group-level $\log_{10}$(odd ratios) of non-synonymous vs. non-coding variants for ALS and FTD cases compared to controls. (b) Individual-level distributions of log10(non-synonymous/non-coding) ratios for ALS, FTD, and controls. Control, n=144; ALS, n=216; FTD, n=78; biological replicates. Data are shown as mean ± 95% CI.

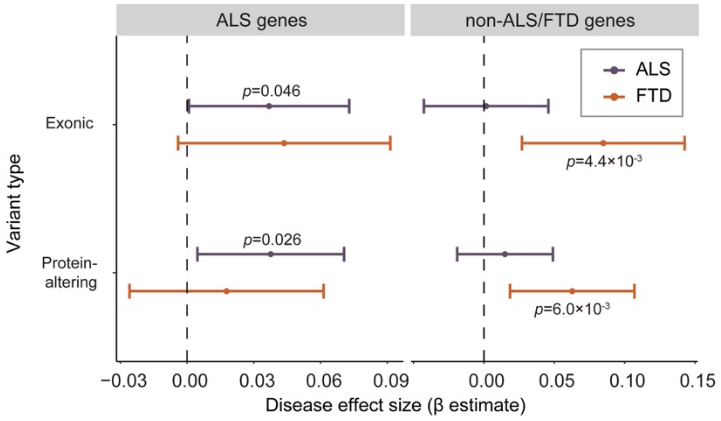


**Extended Data Fig. 5: Enrichment of exonic and protein-altering variants in germline-free ALS and FTD cases.** Somatic variant burdens in ALS genes and non-ALS/FTD neurodegeneration/dementia-related genes were compared between germline-free ALS and FTD cases and normal controls. Enrichment significance and 95% confidence intervals were estimated using a linear mixed model, adjusting for average read-depth, sex, post-mortem interval, sequencing batch, and number of samples per donor as potential confounders. Unadjusted *p* values are shown, as the tests examine biologically distinct but non-independent categories with differing background mutation structures. Control, n=144; ALS, n=216; FTD, n=78; biological replicates.

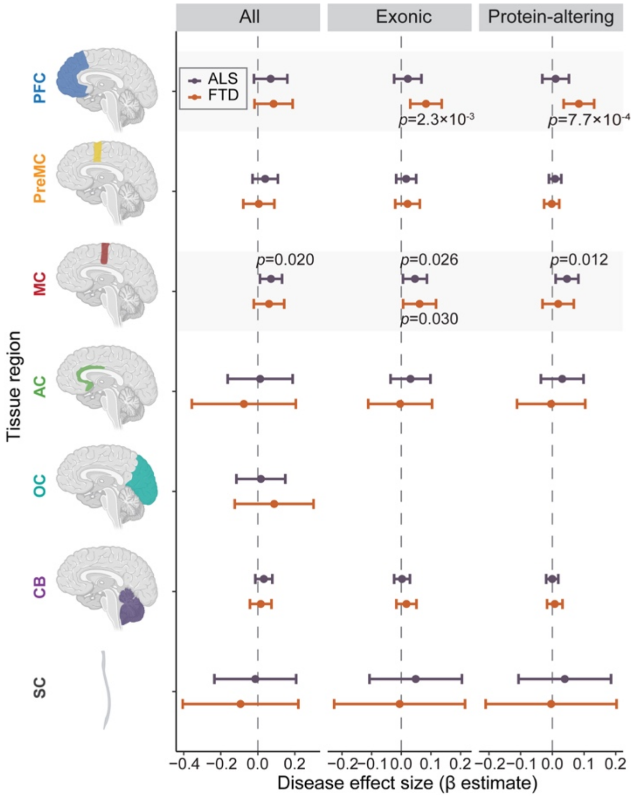


**Extended Data Fig. 6: Enrichment of overall somatic variants across different brain regions of germline-free ALS and FTD cases compared to normal controls.** All somatic variants within the entire set of targeted neurodegenerative genes were analyzed to test for enrichment. The significance of enrichment and 95% CI were estimated while controlling for potential confounding factors including average read-depth, sequencing batch, sampled individuals using a linear mixed model. Unadjusted *p* values are shown, as the tests examine biologically distinct but non-independent categories with differing background mutation structures. Control, n=144; ALS, n=216; FTD, n=78; biological replicates.

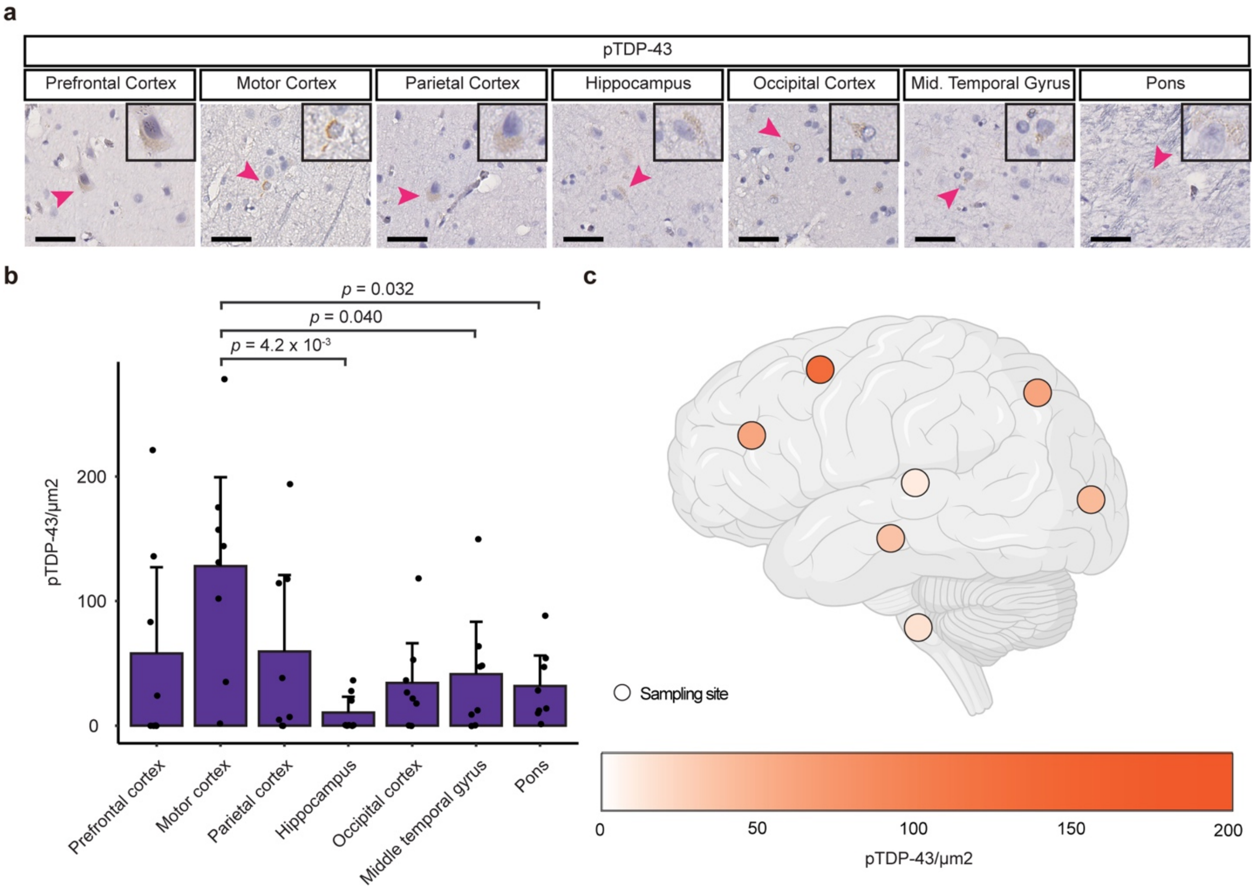


**Extended Data Fig. 7: Quantification of phospho-TDP43 staining in brain tissue sections from the sALS case carrying the *TARDBP* (p.L248F) somatic variant. (a)** Sections of different brain regions of the sALS case stained with a phospho-TDP43 antibody. Scale bar = 40 µm. Arrowheads indicate the cells shown in the insets, which are magnified to twice their original size. (b) pTDP-43 levels across multiple brain regions. Each dot represents the quantification of pTDP-43 in a non-overlapping tissue section. Eight tissue-section-level biological replicates were included per brain region. Differences in pTDP-43 levels across brain regions were evaluated by one-way ANOVA with post hoc Tukey's HSD correction for multiple comparisons. Bar graph, mean ± 95% CI. (c) Regional distribution of mean pTDP-43 levels across brain regions.

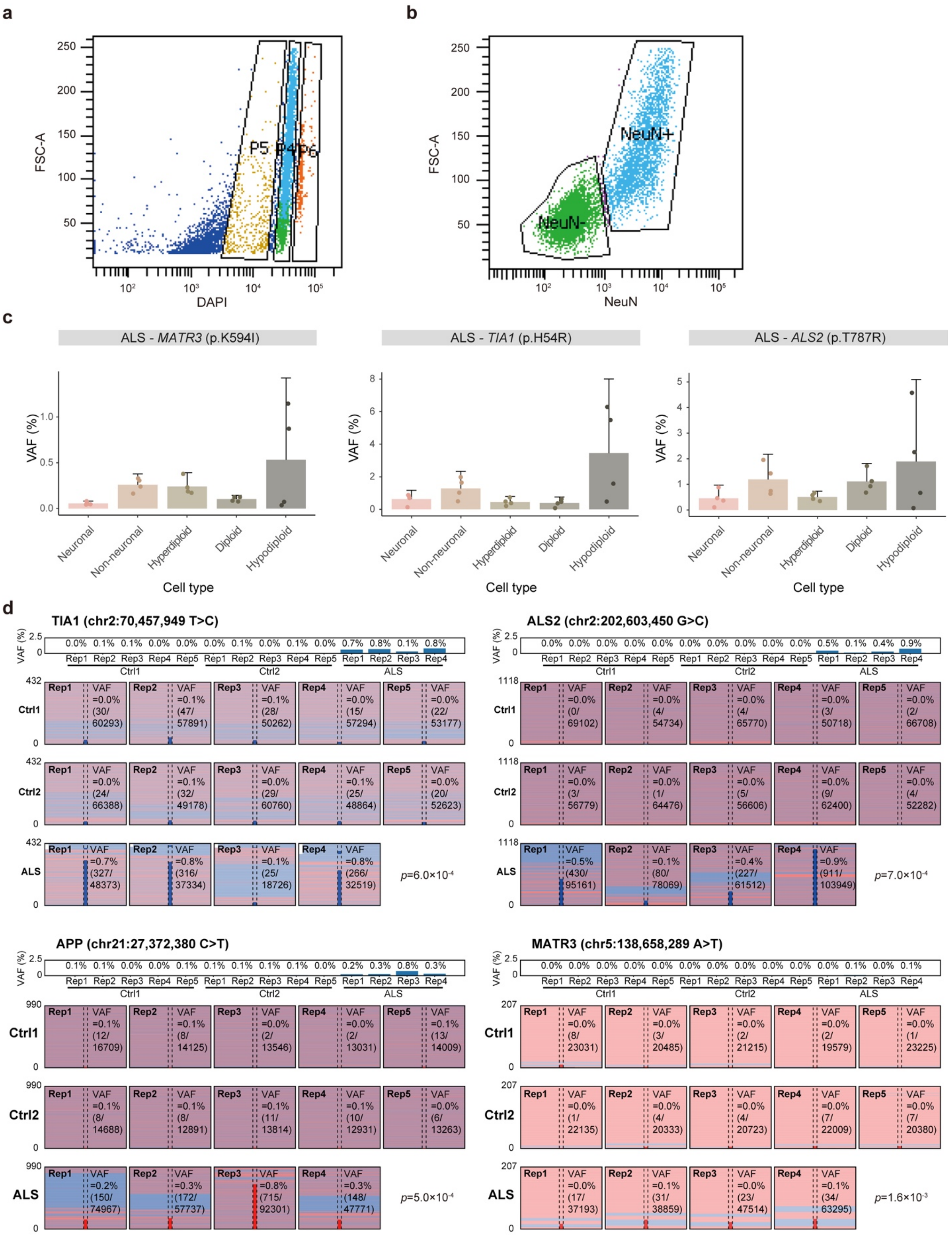
a
FSC-A
DAPI
P5
P4
P6
b
FSC-A
NeuN
NeuN+
NeuN-
c
ALS - MATR3 (p.K594I)
ALS - TIA1 (p.H54R)
ALS - ALS2 (p.T787R)
VAF (%)
Neuronal
Non-neuronal
Hyperdiploid
Diploid
Hypodiploid
Cell type
d
TIA1 (chr2:70,457,949 T>C)
ALS2 (chr2:202,603,450 G>C)
APP (chr21:27,372,380 C>T)
MATR3 (chr5:138,658,289 A>T)
VAF (%)
Rep1
Rep2
Rep3
Rep4
Rep5
Ctrl1
Ctrl2
ALS
p=6.0×10-4
p=7.0×10-4
p=5.0×10-4
p=1.6×10-3

**Extended Data Fig. 8: Predicted deleterious somatic variants are enriched in hypodiploid cells.** All nuclei were stained with DAPI and AF488-conjugated anti-NeuN antibodies to distinguish neuronal and non-neuronal nuclei. (a) Nuclei with diploid DNA content (P4), hypodiploid DNA content (P5), and hyperdiploid DNA content (P6). Cell debris and doublets were removed by prior gates (forward-scatter (FSC-H) versus FSC-A and side-scatter (SSC-H) versus SSH-W). (b) Neuronal and non-neuronal nuclei selected from NeuN staining. Neuronal nuclei typically had larger sizes as indicated by the FSC-A values. (c) VAFs of somatic variants in FANS sorted cell types. Five hundred neuronal (NeuN+), non-neuronal (NeuN-), diploid (DAPI), hyperdiploid (High DAPI) and hypodiploid (Low DAPI) cells were each sorted for amplicon sequencing with four replicates. Bar graph, mean ± 95% CI. (d) Visualization of variant-supporting reads at four somatic variant sites from validation sequencing of sorted neurons. Each subpanel shows a zoomed-in view of variant-supporting reads, with a consistent scale applied per variant. Pink and light blue horizontal lines represent the reads from positive and negative strands, respectively. Empirical two-sided *p*-values were calculated using a group label permutation test (10,000 iterations) comparing the mean VAF between variant carrier and control groups.

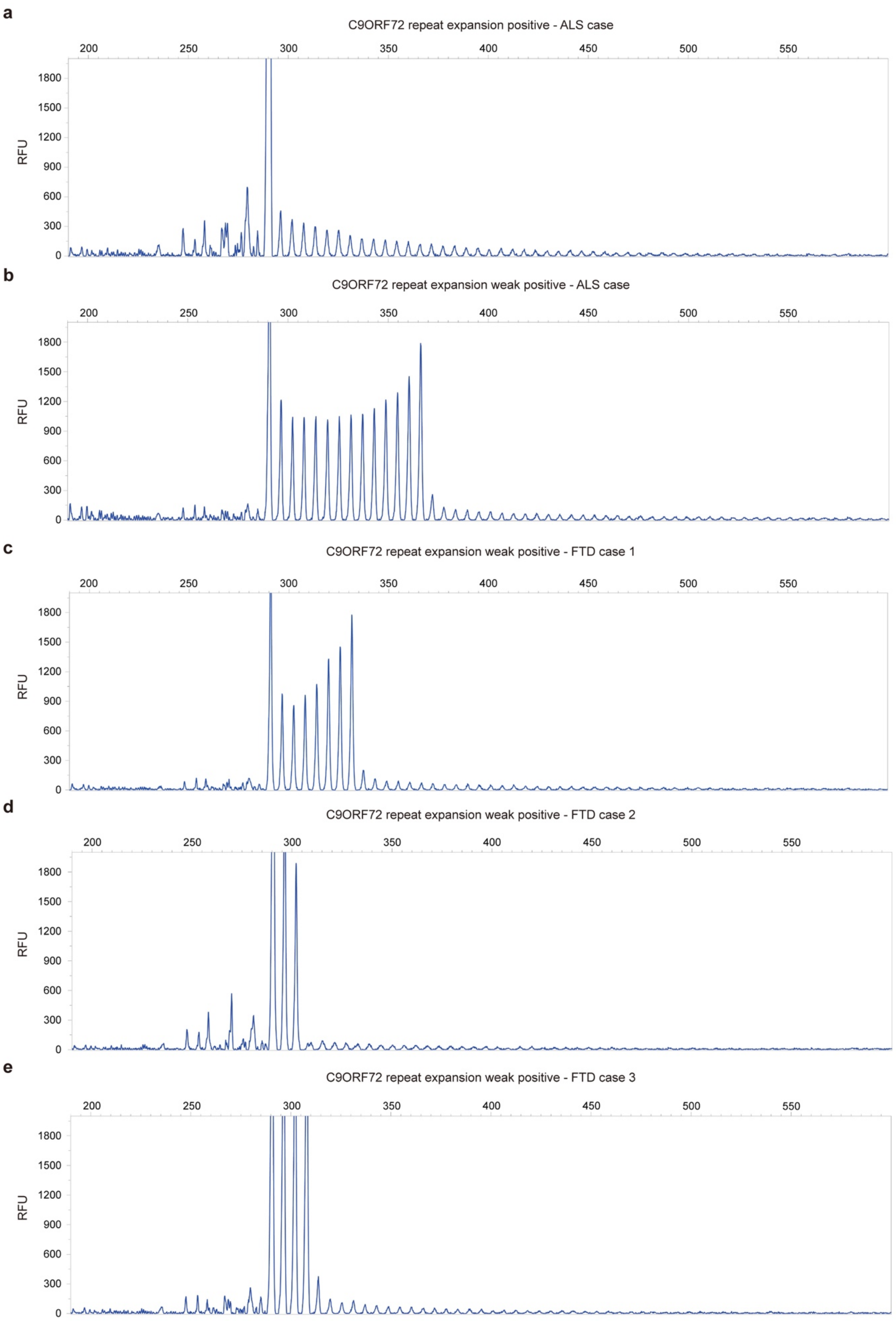
a
C9ORF72 repeat expansion positive - ALS case
b
C9ORF72 repeat expansion weak positive - ALS case
c
C9ORF72 repeat expansion weak positive - FTD case 1
d
C9ORF72 repeat expansion weak positive - FTD case 2
e
C9ORF72 repeat expansion weak positive - FTD case 3
200
250
300
350
400
450
500
550
RFU
0
300
600
900
1200
1500
1800

**Extended Data Fig. 9: Detection of somatic *C9orf72* repeat expansion using repeat-primed PCR.** Electropherograms of cerebellar samples from five ALS and FTD cases. (a) An ALS case exhibited a major peak, indicating a wild-type allele with two repeats. The repeat expansions in this sample showed peaks with high peak heights (reproduced from Extended Data Fig. 2c). (b-e) Four ALS and FTD cases showed major peaks indicating wild-type alleles with various repeat sizes (15, 9, 4, and 5 repeats, respectively). The repeat expansions in these cases showed reduced peak heights compared to the sample shown in (a), suggesting the presence of somatic *C9orf72* repeat expansion. RFU: Relative fluorescence units. X-axis denotes the fragment size (bp).

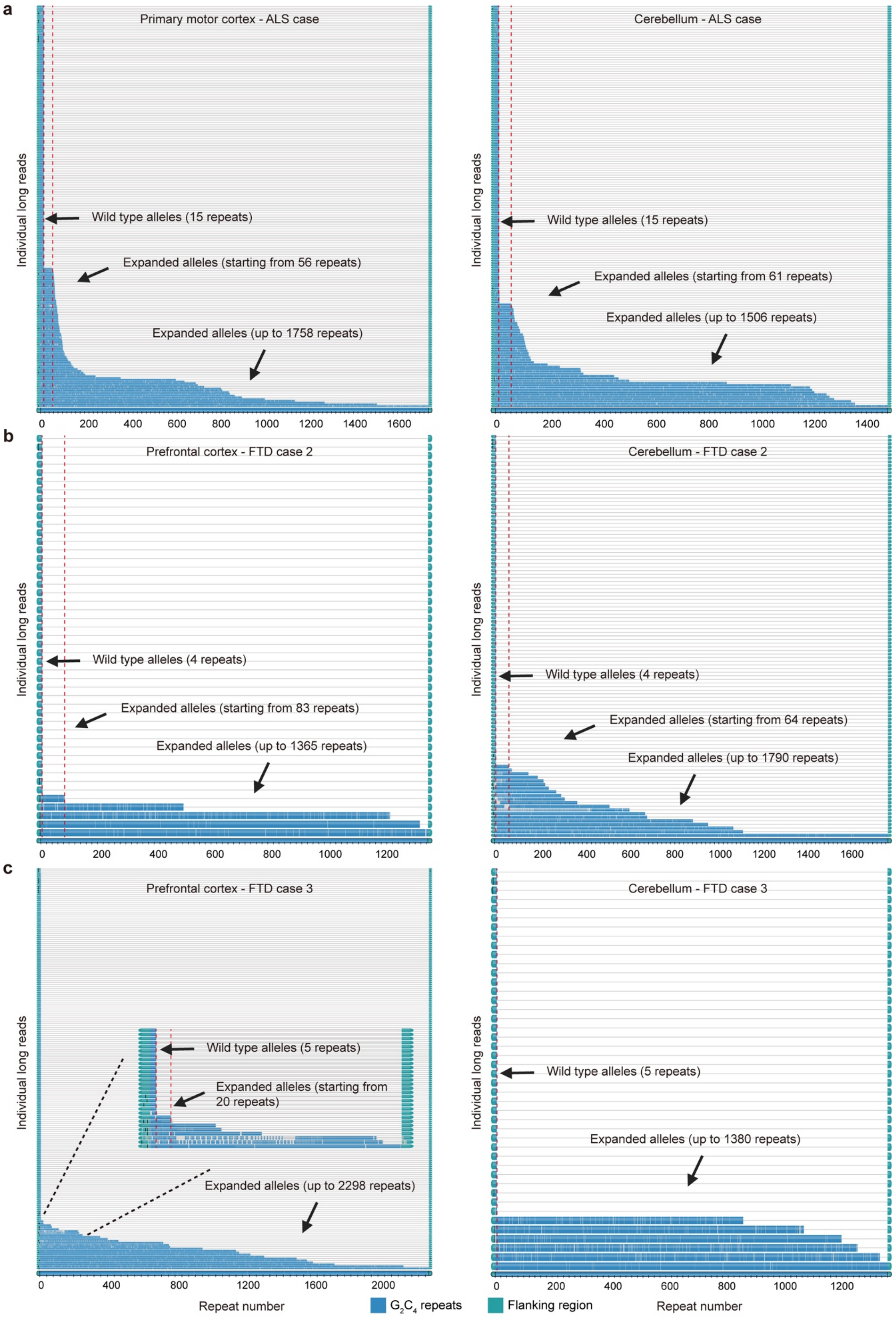
a
Primary motor cortex - ALS case
Wild type alleles (15 repeats)
Expanded alleles (starting from 56 repeats)
Expanded alleles (up to 1758 repeats)
Cerebellum - ALS case
Wild type alleles (15 repeats)
Expanded alleles (starting from 61 repeats)
Expanded alleles (up to 1506 repeats)
b
Prefrontal cortex - FTD case 2
Wild type alleles (4 repeats)
Expanded alleles (starting from 83 repeats)
Expanded alleles (up to 1365 repeats)
Cerebellum - FTD case 2
Wild type alleles (4 repeats)
Expanded alleles (starting from 64 repeats)
Expanded alleles (up to 1790 repeats)
c
Prefrontal cortex - FTD case 3
Wild type alleles (5 repeats)
Expanded alleles (starting from 20 repeats)
Expanded alleles (up to 2298 repeats)
Cerebellum - FTD case 3
Wild type alleles (5 repeats)
Expanded alleles (up to 1380 repeats)
Individual long reads
Repeat number
G2C4 repeats
Flanking region

**Extended Data Fig. 10: Detection of somatic *C9orf72* repeat expansion using targeted long-read sequencing.** Waterfall plots from targeted long-read sequencing results of brain tissues from ALS and FTD cases revealed the presence of both wild-type alleles and expanded alleles. (a) In an ALS case, the primary motor cortex and cerebellum samples showed a wild-type allele with 15 repeats and expanded alleles raging from 56 to 1758 repeats. (b) In an FTD case, the prefrontal cortex and cerebellum samples showed a wild-type allele with 4 repeats and expanded alleles raging from 64 to 1790 repeats. (c) In an FTD case, the prefrontal cortex and cerebellum samples showed a wild-type allele with 5 repeats and expanded alleles raging from 20 to 2298 repeats. Red dashed lines indicate the sizes of the wild-type alleles and the shortest expanded alleles. Each row in the waterfall plots represents an individual sequencing read, with flanking regions shown in green and the *C9orf72* repeat expansions in blue. The x-axis indicates the number of repeats.

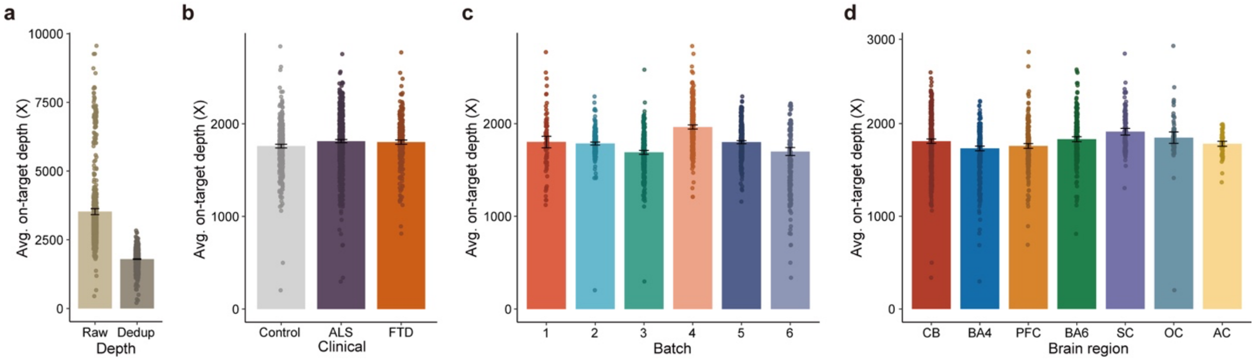


**Supplementary Fig. 1: Depth distribution of MIP panel sequencing data.** Average sequencing depth of the data is depicted across different conditions: (a) before and after UMI deduplication (n=1,787), (b) across clinical conditions (n=516, 938, and 375, respectively; 42 samples are included in both ALS and FTD as they originate from ALS-FTD patients), (c) by sequencing batch (n=91, 307, 358, 378, 378, and 275, respectively), and (d) across brain regions (n=529, 393, 295, 290, 132, 75, and 73, respectively). All samples represent biological replicates. Bar graph, mean ± 95% CI.

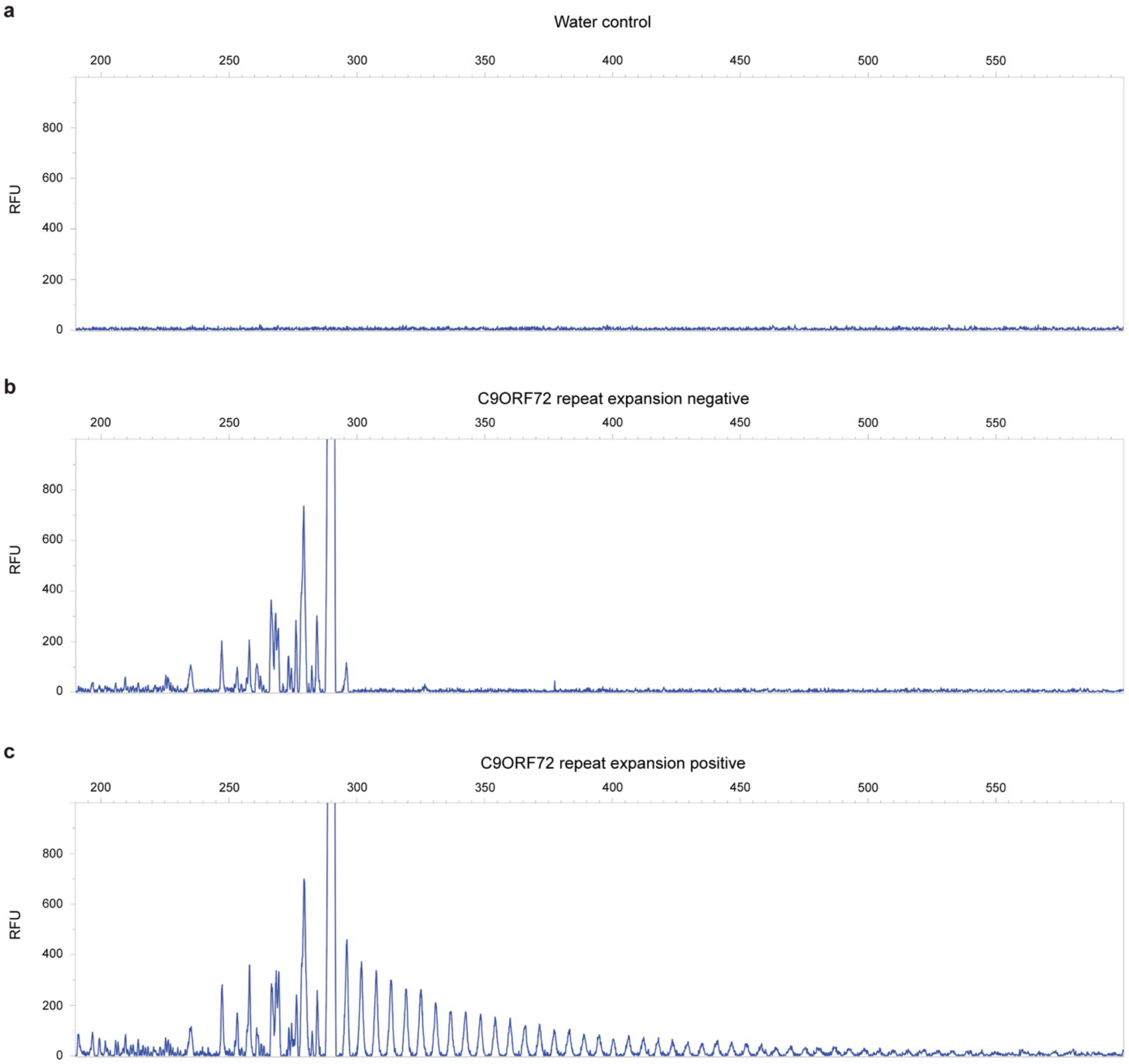


**Supplementary Fig. 2: Genotyping of *C9ORF72* hexanucleotide repeat expansions for ALS and FTD cases using repeat-primed PCR assays.** The presence of *C9ORF72* hexanucleotide repeat expansions in 291 ALS and 117 FTD cases was assessed by repeat-primed PCR assays. (a) The electropherogram of a water control without *C9ORF72* hexanucleotide repeat expansions. (b) A negative sample with a low number of *C9ORF72* hexanucleotide repeat expansions. (c) A positive sample with a large number of *C9ORF72* hexanucleotide repeat expansions. RFU: Relative fluorescence units. X-axis denotes the fragment size (bp).

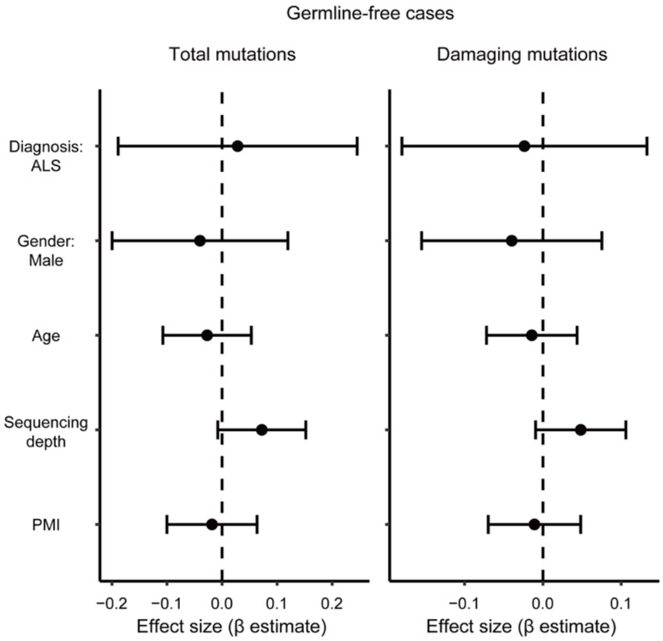


**Supplementary Fig. 3: Analysis of somatic SNVs in bulk RNA-seq data of brain and spinal cord tissues of sALS cases detected by RNA-MosaicHunter.** Linear regression modeling did not find a significant enrichment of somatic SNVs in sALS cases that lack pathogenic germline variants in ALS genes. PMI, post-mortem interval. Bulk RNA-seq data from 682 tissues of 143 sALS cases and 107 tissues of 23 control cases were analyzed. The significance of enrichment and 95% CI were estimated while controlling for potential confounding factors including gender, age, and sequencing and sample qualities using a linear mixed model.

### Supplementary Note

#### MIP panel design and sequencing of postmortem tissues from diseased and control individuals

We designed a MIP panel targeting the exons and exon-intron junctions of 88 neurodegeneration/dementia-related genes, which included 34 ALS/FTD genes, 10 Alzheimer's disease genes, 28 Parkinson's disease genes, and 16 genes associated with other rare neurodegenerative disorders (Supplementary Table 2). We performed MIP panel sequencing at ~1,800X average sequencing depth after deduplication based on unique molecular identifiers (UMIs) (Fig. 1b and Supplementary Fig. 1), with a similar distribution of sequencing depth across batches, disease conditions, and tissue regions (Supplementary Fig. 1). The variance of depth, along with the batch and sample information, were considered as covariates in the variant burden test. A total of 938, 375, and 516 samples from 291 ALS, 117 FTD, and 144 neurotypical control individuals respectively were sequenced (Fig. 1a, 1c and Supplementary Table 1). Of the ALS and FTD cases, nine were diagnosed with both ALS and FTD. Therefore, 42 samples from these cases were included in both conditions, leading to a total of 1,787 unique samples.

#### Benchmarking of the custom pipeline for somatic variant calling

We performed spike-in experiments by mixing two human samples from the Genome in a Bottle Consortium (GIAB) at VAFs of 10%, 5%, 2.5%, 1%, and 0.5% and estimated theoretical sensitivity across sequencing depths (Extended Data Fig. 1a, b; see Methods). Double-called variants identified by Mutect2 and Pisces were excluded from the final call set due to high false positive and low validation rates (Extended Data Fig. 1c, d). High sensitivity and precision were achieved for the remaining Replow-based double-called variants (Replow-Mutect2 and Replow-Pisces) while maintaining a low false positive rate across the low VAFs compared to the somatic variants called by each caller. The MIP sequencing and our custom pipeline together allowed us to confidently identify somatic variants with a low false positive rate at VAF as low as 0.5%. The observed VAFs of somatic variants were well in line with the target VAFs at all five VAF levels.

#### Detection of somatic variant candidates arising from sample contamination

We identified low-level DNA contamination derived from another sample in 29 out of 1,787 samples (13 ALS, 10 FTD, and 6 control samples). Germline variants from the contaminant mimicked low-VAF somatic variants, leading to false positive calls. To address this, we implemented a module to identify low-level contamination and filter out candidates originating from the contaminant. By comparing the somatic candidate set of a given sample with the germline call set of other individuals, sample contamination was determined if the sample had $\geq$ 40 low-VAF somatic candidates that were also observed as germline variants in another specific individual. In such cases, the germline variants of the matched individual were considered as potential sources of false positive calls, and all matching somatic candidates of the contaminated sample were filtered out. After this filtration, six of the 29 samples still harbored a total of seven somatic variant candidates. To ensure these were not artifacts of contamination, the remaining seven candidates from the contaminated samples were reexamined to verify the absence of corresponding variants within the contaminant sources, confirming these as independent candidates.

**Hypodiploid nuclei indicate apoptotic cells**

Previous flow cytometric studies have established that reduced DNA content, leading to a hypodiploid cell population, is a hallmark of late-stage apoptosis, primarily due to leakage of endonuclease-cleaved DNA fragments. These fragmented cells form a distinct hypodiploid population, often referred to as the "sub-G1" peak.

**Burden analysis of somatic variants using linear mixed model**

For both MIP and RNA sequencing data, linear mixed-effect regression models (linear mixed models) were used to evaluate the relationships between somatic variant burden and clinical conditions, while accounting for other covariates that may affect the burden. An individual-level analysis framework was used, in which candidate somatic variants were aggregated across all tissue samples from each individual. Individual-wise variant lists were constructed by merging all candidate variants from the same donor, with overlapping variants across multiple tissue samples counted only once to avoid overestimation. A linear mixed-effect regression model was then applied to evaluate associations between somatic variant burden and clinical conditions, while adjusting for potential covariates. Clinical conditions and covariates of interest—including sex, postmortem interval (PMI), average sequencing depth, and the number of samples per donor—were modeled as fixed effects. Sequencing batch (batch 1 to 6) was modeled as a random effect to account for variation introduced by sample clustering within the same batch.

The somatic variant burden per individual was modeled as: $y_{ij}=\mu+\alpha_i+\beta_i+\gamma_i+\delta_i+n_i+U_i+\varepsilon_i$, where $y_i$ is the somatic variant burden of donor $i$ (normalized per megabase), $\mu$ is the average variant burden in the normal condition, $\alpha_i$ is the fixed effect of PMI, $\beta_i$ is the fixed effect of disease status (ALS or FTD) compared to normal controls, $\gamma_i$ is the fixed effect of sex, $\delta_i$ is the fixed effect of average sequencing depth across all samples from donor $i$, and $n_i$ is the fixed effect accounting for the number of samples per donor. $U_{ij}\sim N(0, \sigma_r^2)$ represents the random effect of sequencing batch, and $\varepsilon_i\sim N(0, \sigma^2)$ is the residual error term. A covariate with a p-value < 0.05 was considered to be significant, based on a t-test using the Satterthwaite approximation of degrees of freedom. To test the burden of somatic variants in different genomic regions, a linear mixed model was fitted to the corresponding variant counts of specific type (e.g. exonic). To test the burden of somatic variants in different brain regions, samples were first divided by the sequenced region and then a linear mixed model was fitted for each region group. All models were fitted using R (v4.1.0) with the lme4 (v1.1.30) and the lmerTest (v3.1.3) R packages.
